# A quiescent state following mild sensory arousal in *Caenorhabditis elegans* is potentiated by stress

**DOI:** 10.1101/784306

**Authors:** Patrick D. McClanahan, Jessica M. Dubuque, Daphne Kontogiorgos-Heintz, Ben F. Habermeyer, Joyce H. Xu, Anthony M. Ma, David M. Raizen, Christopher Fang-Yen

## Abstract

An animal’s behavioral and physiological response to stressors includes changes to its responses to stimuli. How such changes occur is not well understood. Here we describe a *Caenorhabditis elegans* quiescent behavior, post-response quiescence (PRQ), which is modulated by the *C. elegans* response to cellular stressors. Following an aversive mechanical or blue light stimulus, worms respond first by briefly moving, and then become more quiescent for a period lasting tens of seconds. PRQ occurs at low frequency in unstressed animals, but is more frequent in animals that have experienced cellular stress due to ultraviolet light exposure as well as in animals following overexpression of epidermal growth factor (EGF). PRQ requires the function of the carboxypeptidase EGL-21 and the calcium-activated protein for secretion (CAPS) UNC-31, suggesting it has a neuropeptidergic mechanism. Although PRQ requires the sleep-promoting neurons RIS and ALA, it is not accompanied by decreased arousability, and does not appear to be homeostatically regulated, suggesting that it is not a sleep state. PRQ represents a simple, tractable model for studying how neuromodulatory states like stress alter behavioral responses to stimuli.

## Introduction

Animals exhibit behavioral and physiological responses to stressors such as adverse environmental conditions, food scarcity, and predator threat. One of the ways in which stress affects behavior is by modulating behavioral responses to stimuli. For example, in rats, stress increases sensitivity to auditory stimuli^1^. Acute stress causes inhibition of responses to food stimuli^2^ and prolonged stress decreases responses to social cues^3^. In mammals, stress hormones like cortisol upregulate behavioral responses to predators, including quiescent states of defensive freezing^4^ and tonic immobility^5^.

In humans, severe stress can induce the maladaptive responses to stimuli observed in post-traumatic stress disorder^6^, in addition to increased risks of anxiety, depression, and other disorders^7^. Despite its significant medical and public health implications, our understanding of the regulation of stimulus response behavior by stress remains quite limited^8^.

The study of genetically tractable models can improve our understanding of the regulation of behavior under stress. The roundworm *Caenorhabditis elegans* exhibits a stereotypical response to many stressors. Following exposure to a stressor like heat or UV radiation, the worm exhibits a period of quiescence^9^ that aids survival^10–12^. This quiescence is mediated by the ALA and RIS neurons, which can be activated by the EGF homologue LIN-3^12^, releasing a cocktail of neuropeptides, including those encoded by *flp-13*^13,14^. Like sleep, this recovery quiescence is associated with reduced responsiveness to some stimuli^10,15^, but the quiescence can be rapidly reversed by a sufficiently strong stimulus^10^. Quiescence after stress is therefore similar to recovery sleep or quiescence in other animals and is referred to as stress-induced sleep (SIS)^16^. In mammals, EGF signaling promotes behavioral quiescence and sleep^17,18^ and is released following stress^19^, suggesting a conserved role for EGF in animal stress response regulation.

Like its physiological stress response, the *C. elegans* acute escape behavior has been extensively characterized. Six touch-receptor neurons sense gentle mechanical stimuli to the body as well as substrate vibration, allowing the animal to quickly accelerate away from the stimulus^20,21^. Similar escape responses occur following aversive chemical^22^, osmotic^23^, optical^24^, and thermal^25^ stimuli. In invertebrates, tyramine and octopamine function analogously to epinephrine and norepinephrine in the mammalian fight or flight response^26,27^, and the *C. elegans* locomotor escape response is accompanied by a tyramine-dependent^27^ inhibition of lateral head movements^20^. Mutations that impair touch sensitivity or head movement suppression during backing increase the worm’s susceptibility to predation by trap-forming *Drechslerella doedycoides* fungi^28^. However, little is known about how stress affects *C. elegans* escape behavior.

Here we describe a freezing behavior following the locomotor escape response to mild aversive stimuli. We show that this behavior, which we call post response quiescence (PRQ), is enhanced following the EGF-mediated cellular stress response. We find that some of the genes and neurons required for SIS are also required for PRQ. Like SIS, PRQ requires genes involved in the processing and secretion of neuropeptides, and the sleep active interneurons RIS and ALA. Despite its genetic and anatomical overlap with *C. elegans* sleep states, PRQ appears to lack two of the behavioral characteristics of sleep – decreased arousability and homeostatic regulation – suggesting it is distinct from sleep. The occurrence of PRQ during an escape response and upregulation by the stress response pathway suggest a role in predator evasion similar to freezing behavior in mammals.

## Results

### Quiescence increases after mechanosensory response during UV stress-induced sleep

We asked how cellular stress affects the *C. elegans* behavioral response to aversive mechanical stimuli. To induce SIS, we exposed young adult wild-type (WT) animals to UV radiation^29^. After the given exposure to UV radiation, the animals became sterile and died within one week **(Supplementary Fig. S1)**, suggesting that they sustain considerable damage^29^. To track behavior, we imaged worms isolated in wells of a multi-well device (WorMotel)^30^ and used frame subtraction^31^ to measure behavioral activity and quiescence (see Methods). Consistent with previous reports, quiescence reached a maximum 1 – 2 hours following UV-C exposure, then decreased over the next 5 hours **(Fig. 1a)**. The peak fraction of time spent quiescent was about 70%, allowing for the detection of both increases and decreases in quiescence.

**Figure 1.**
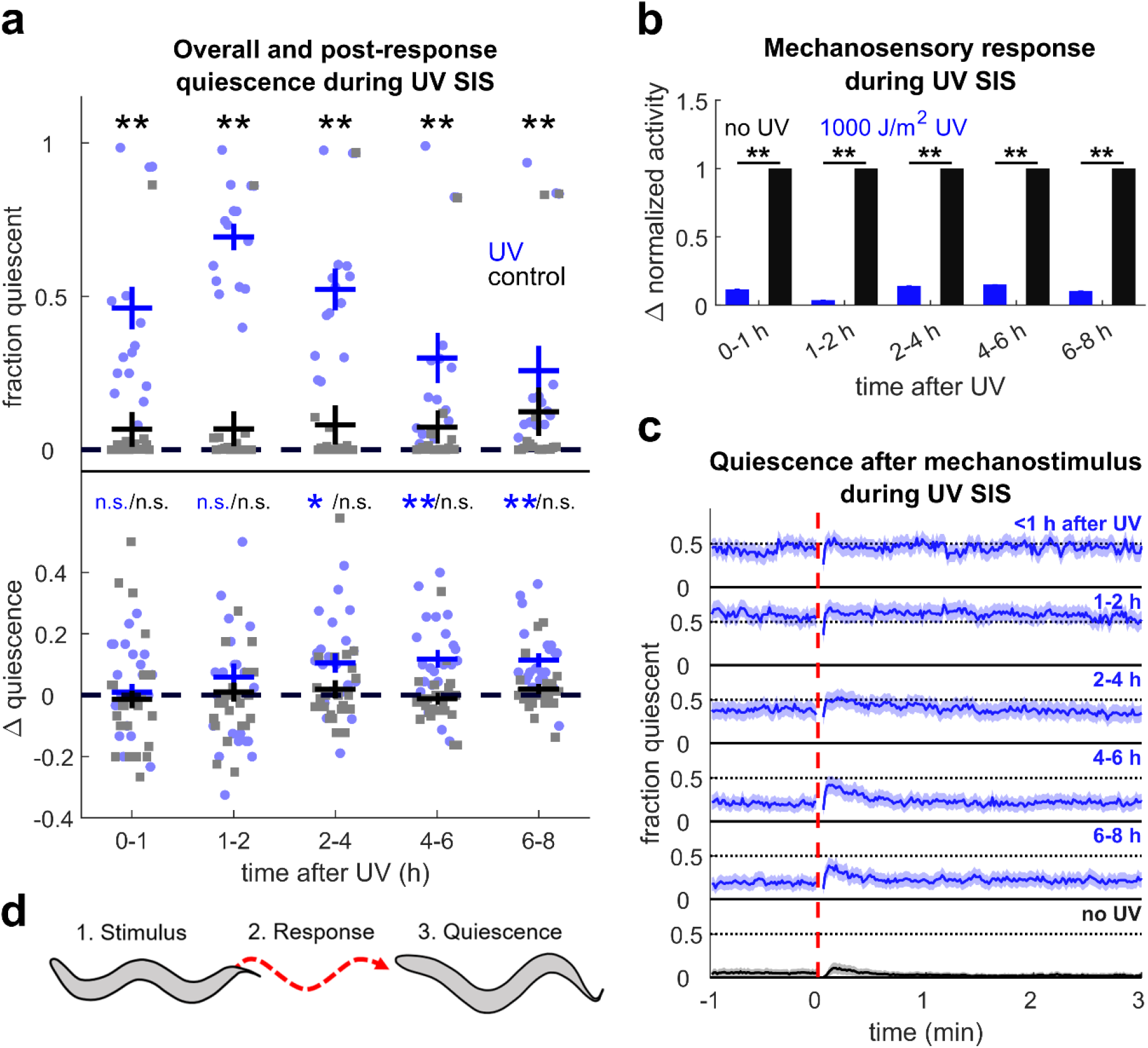
Mechanosensory stimulus during UV SIS elicits post response quiescence. **(a) (top)** Fraction of time quiescent of WT animals after 1000 J/m^2^ UV (blue) and non-exposed controls (black) in the absence of mechanostimuli. Dots represent individual animal quiescence in 1 or 2 hour bins. n = 15 animals per condition, four replicates. ** denotes p < 0.01 comparing UV-treated to controls (Wilcoxon rank-sum test with Bonferroni correction for five comparisons) **(bottom)** Quantification of PRQ. Change fraction in quiescent (Δ quiescence) is the peak quiescence within two minutes after mechanostimulus minus the peak quiescence within two minutes before mechanostimulus in the time bin specified. Typically, four or eight responses were averaged per animal and smoothed with a 10 s averaging filter. n = 22 animals per condition, two replicates. ** denotes p < 0.01, * denotes p < 0.05 (two-sided Wilcoxon signed rank test with Bonferroni correction for five comparisons). Error bars indicate ± SEM. **(b)** Mechanosensory response during UV SIS normalized to non-UV controls. Response is the average activity from the five seconds after the stimulus minus the average activity in the minute before the stimulus. The data are from the same animals as panel a (bottom). ** denotes significance (two-sided Wilcoxon rank-sum test with α = 0.01 and Bonferroni correction for five comparisons). Error bars indicate SEM. **(c)** Fraction quiescent before and after stimulus (dashed red line) 0-1, 1-2, 2-4, 4-6, and 6-8 h (blue) after UV treatment and in non-UV controls averaged throughout the experiment (black). Data are from the same animals as panel a (bottom) and panel b. Shading denotes ± SEM. **(d)** Schematic of post-response quiescence: (1) following stimulus, (2) the animal responds by moving, and (3) subsequently becomes quiescent.

To deliver mechanical stimuli, we attached a WorMotel containing the worms to an audio loudspeaker, which was driven by a computer and amplifier to produce 1 s duration, 1 kHz frequency vibrations every 15 minutes. Substrate vibrations elicit a mechanosensory response mediated by the gentle touch receptors^32,33^ without considerably altering overall the time course of SIS **(Supplementary Discussion 1, Supplementary Fig. S2)**. Both chemosensory and photosensory responsiveness are known to decrease during SIS^10,29^, and mechanosensory arousability decreases in another *C. elegans* sleep state, developmental lethargus^34,35^. Consistent with decreased overall sensory arousability, mechanosensory responses decreased during SIS relative to controls, displaying a minimum at the time of peak quiescence **(Fig. 1b)**.

Despite an overall decrease in activity during SIS, most worms usually showed movement in response to the mechanical stimulus. We noticed that UV-treated worms sometimes froze for short periods (tens of seconds) following their brief locomotor response **(Supplementary Vid. 1)**. Control animals, which had not been exposed to UV light, rarely displayed this freezing behavior, instead displaying normal mechanosensory responses **(Supplementary Vid. 2)**.

To quantify this freezing behavior, we compared the peak fraction of animals quiescent before and after the stimulus (see Methods). Significant increases in quiescence following mechanostimulus occurred after the time of deepest SIS, becoming statistically significant in the 2-4, 4-6, and 6-8 hour periods (p = 0.032, 0.007, and 0.001, respectively) **(Fig. 1a,c)**. We did not find a significant increase in quiescence in the first and second hours after UV exposure, or in control animals at any time **(Fig. 1a)**, although we occasionally observed similar behavior in untreated controls **(Supplementary Vid. 3)**. We refer to this behavior, a cessation of body movement following a mechanosensory response, as post-response quiescence (PRQ) **(Fig. 1d)**.

Because SIS consists of alternating bouts of activity and quiescence, it is possible that PRQ results from the synchronization of quiescent bouts after a mechanosensory response. To test this hypothesis, we simulated the synchronization of quiescent bouts by temporally aligning baseline (pre-stimulus) SIS data from experimental animals according to the beginnings of active or quiescent bouts and compared the resulting traces to PRQ in the same animals. Although aligning quiescent bouts does produce a peak in quiescence, the peak resulting from the alignment of quiescent bouts has a much smaller area than that observed during PRQ, implying that PRQ cannot result from the temporal synchronization of the normal quiescent behavior in UV SIS or after EGF overexpression (discussed below) **(Supplementary Discussion 2, Supplementary Fig. S4)**.

Together, these results suggest that UV stress affects mechanosensory response behavior in two ways. First, the overall locomotor activity following the stimulus is reduced, consistent with a lower arousability during SIS. Second, beginning about two hours after UV exposure, the animals freeze after initiation of an otherwise normal mechanosensory response, and before returning to baseline quiescence levels.

### Post response quiescence is enhanced following LIN-3C / EGF overexpression

SIS caused by UV and other stressors occurs following activation of the ALA neuron, and possibly the RIS neuron^12^. These neurons both express LET-23, the receptor for the EGF homologue LIN-3. Both *lin-3* and *let-23* hypomorphs show a reduction in SIS after heat stress, and LIN-3 (EGF) overexpression induces quiescence resembling SIS. These findings suggest that EGF plays a key role in the induction of SIS^9,10^. We therefore asked whether PRQ is regulated downstream of EGF, or is part of an EGF-independent UV stress response. To address this question, we induced overexpression of EGF by heating (32 °C for 10 min) animals carrying a *lin-3c* transgene under the control of a heat shock promoter. In contrast to reported severe heat shock (35-37 °C for 30 min^10^), this mild heat shock paradigm had minimal effects on control animals that did not carry the hs:EGF transgene. As reported^9,29^, EGF overexpression caused an overall increase in quiescence and decrease in arousability with dynamics similar to that of UV SIS **(Fig. 2a,b)**. We then used our mechanostimulus protocol to study the mechanosensory response.

**Figure 2.**
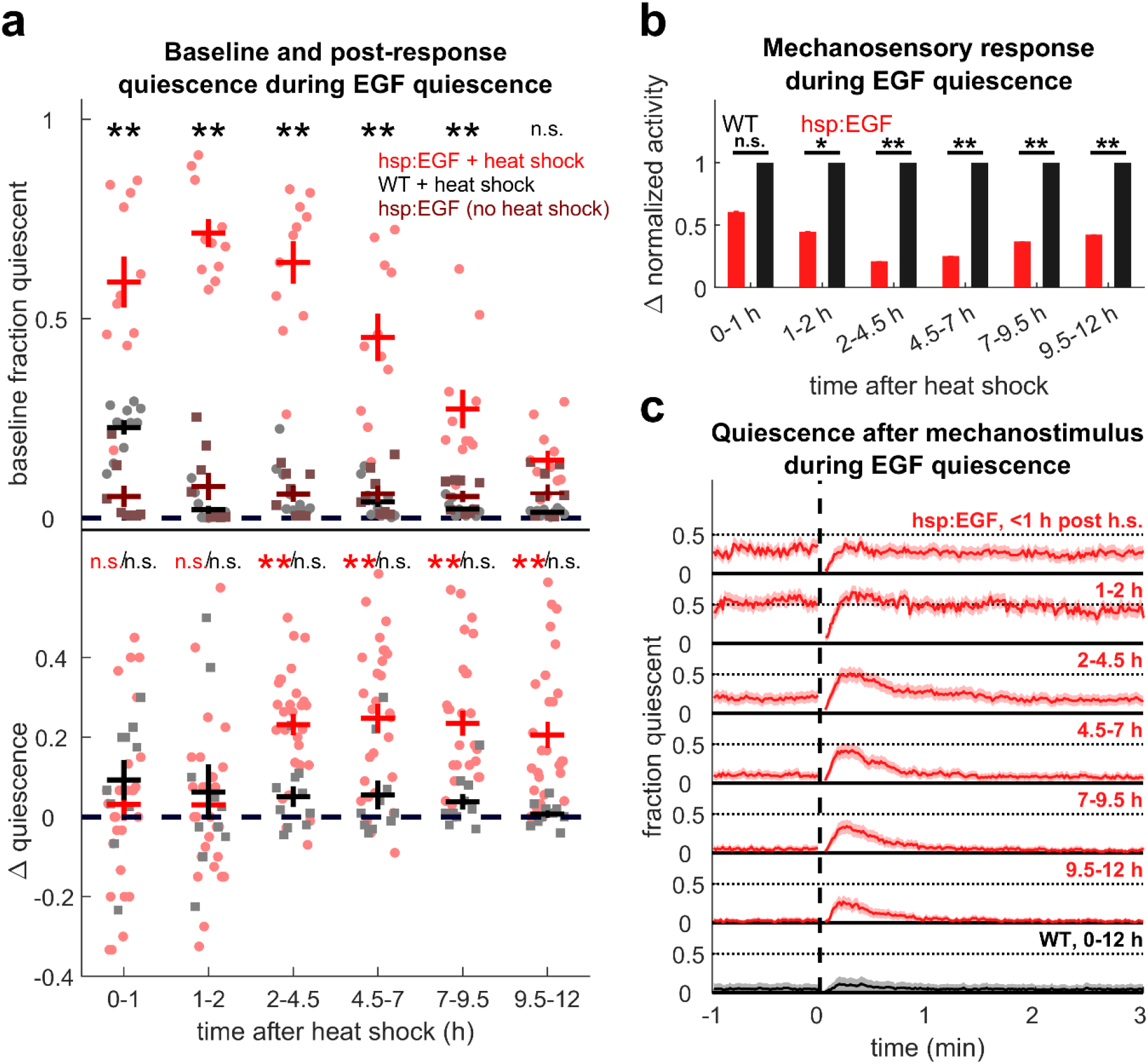
PRQ occurs after EGF overexpression. **(a) (top)** Fraction of time quiescent of hsp:EGF (red, circles) and WT (black, circles) after a mild, 10 min, 32 °C, heat shock and hsp:EGF held at room temperature (burgundy, squares) in the absence of mechanostimulation. n = 8 to 11 animals per condition, two replicates. ** denotes p < 0.01 comparing heat shocked hsp:EGF to WT controls (two-sided Wilcoxon rank-sum test with Bonferroni correction for six comparisons) **(bottom)** Quantification of PRQ in hsp:EGF (red) and WT controls (black) after mild heat shock. Δ quiescence was calculated as in Fig. 1a. Pale markers represent individual animal averages. ** denotes p < 0.01, two-sided Wilcoxon signed rank test with Bonferroni correction for six comparisons. Error bars indicate ± SEM. **(b)** Mechanosensory response after EGF overexpression normalized to mildly heat-shocked WT controls. Response is the average activity from the five seconds after the stimulus minus the average activity in the minute before the stimulus. n = 28, four replicates for hsp:EGF and n = 10, two replicates for WT controls. * denotes p < 0.05, ** denotes p < .01, two-sided Wilcoxon ranksum test with Bonferroni correction for six comparisons. Error bars indicate SEM. **(c)** Fraction quiescent before and after stimulus (dashed red vertical line) in hsp:EGF animals 0-1 h, 1-2 h, 2-4.5 h, 4.5-7 h, 7-9.5 h, and 9.5-12 h after heat shock (red) and WT animals 0-12 h after heat shock (black). Shading denotes ± SEM. The animals are the same as those in panel b.

We found that PRQ occurs following EGF overexpression-induced quiescence, and resembles PRQ after UV exposure **(Fig. 2a,c)**. This result is consistent with PRQ being upregulated downstream of or in parallel to EGF signaling during SIS. For our parameters for UV exposure and EGF overexpression, the increase in fraction quiescent following mechanostimulus was greater after EGF overexpression (reaching a maximum of 0.25 ± 0.07 in hours 4.5-7) than after UV exposure (reaching a maximum of 0.12 ± 0.06 in hours 4-6 after UV-C exposure **(Fig. 1a, 2a)**.

### Post-response quiescence following mechanostimulus requires touch receptor neuron function

Though we expected that our mechanical vibration stimulus generated a response and PRQ through the touch receptor neurons, it remained possible that vibration induced PRQ by a mechanism independent of these cells. To ascertain whether PRQ requires the function of the touch receptor neurons, we assayed for PRQ in EGF-overexpressing worms containing a *mec-4* mutation, which causes a defective behavioral response to vibration^20^. We found that *mec-4* mutants show slightly higher quiescence levels than WT controls following EGF overexpression, even in the absence of deliberate mechanostimulation, although mechanostimulation may further reduce WT quiescence levels compared to *mec-4*. This is consistent with observations that this mutant is generally lethargic^20,28,36^ **(Supplementary Fig. S5a,b)**. After mechanostimulus, EGF-overexpressing *mec-4* mutants displayed neither a movement response nor a PRQ response to the vibration **(Supplementary Fig. S5c,d)**. When we restricted analysis to compare the two strains’ PRQ for time periods in which the baseline quiescence level was similar, we found they still lack PRQ, indicating that our failure to detect PRQ in this strain was not due to a quiescence ceiling effect **(Supplementary Fig. S5e,f)**. These results indicate that PRQ occurs downstream of activation of the touch receptor neurons and possibly the escape response that they mediate.

### The propensity and timing of PRQ vary widely after different cellular stressors

In addition to UV radiation, exposure to ethanol, high salt, Cry5b pore-forming toxin, heat shock, and cold shock also induce SIS^10^. To test whether these stressors also promote PRQ, we assessed quiescence changes following substrate vibration in animals exposed to these stressors for 12 h following exposure. We compared the behavioral response of stressed animals and EGF-overexpressing positive controls to negative control animals on the same WorMotel chip (see Methods). Consistent with previous reports^10^, we observed an overall increase in quiescence consistent with SIS in animals exposed to these stressors, although the amount and time course of quiescence varied **(Supplementary Fig. S6a)**.

Because the time course of SIS varies by stressor, and therefore PRQ regulation may also vary, we looked for PRQ following stimuli in 3 h bins throughout the 12 h recording period. In all experiments, EGF-overexpression caused a significant increase in PRQ starting at 3 h. However, to our surprise, we found only one case where PRQ was significantly higher in SIS animals than in control animals that had not been exposed to the stressors: 9-12 h after Cry5b exposure **(Fig. 3, Supplementary Fig. S6b)**. We also observed a non-significant trend toward PRQ 3-9 h after NaCl exposure and after heat shock.

**Figure 3.**
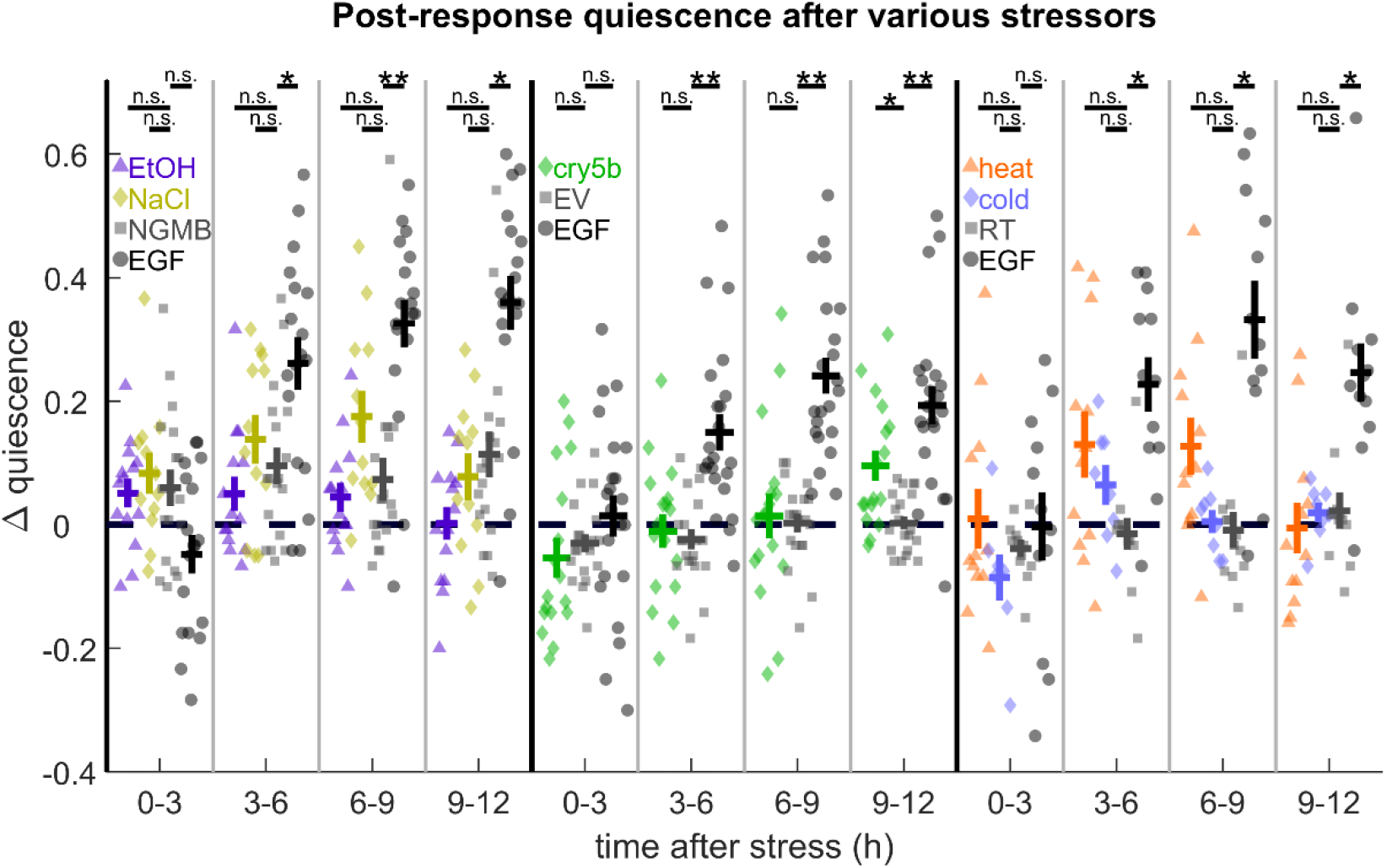
PRQ assays during ethanol, high salt, Cry5b toxin, heat, and cold – induced SIS. PRQ following mechanostimulus after various stressors and EGF overexpression (black circles) compared to experiment-specific negative controls (gray, squares). **(left)** Comparison of PRQ after 15 min immersion in 5% ethanol by volume (purple, triangles), +500 mM NaCl in NGMB (yellow, diamonds), and EGF overexpression (black, circles) to NGMB-immersed controls (gray, squared). **(middle)** Comparison of PRQ after 15 min exposure to Cry5b-expressing bacteria (green, diamonds) or EGF overexpression (black, circles) to empty vector (EV) bacteria-exposed animals (gray, squares). **(right)** Comparison of PRQ after 30 min, 36°C heat shock (orange, triangles) or 24 h, 4°C cold shock (blue, diamonds) to room temperature controls (gray, squares). Pale markers represent individual animal averages over the specified time period. Δ quiescence was calculated as in Fig. 1a. Error bars indicate ± SEM. * denotes p < 0.05 and ** denotes p < 0.01 in a two-sided Wilcoxon rank-sum test after Bonferroni correction for twelve (liquids, left; temperature, right) or eight (Cry5b toxin, middle) comparisons.

We considered the possibility that PRQ requires the dose of the stressor to be within a certain range, and that the stressors we applied were either too weak or too strong to cause PRQ. The dosage of different stressors can be compared via the baseline quiescence they induce. To explore this idea further, we applied our mechanosensory protocol to assay for PRQ in animals exposed to a 36°C heat shock for 15 min or 45 min, which causes lower and higher baseline quiescence, respectively, than in our EGF overexpression protocol. We chose heat shock because the evidence that heat shock SIS is directly caused by EGF signaling is stronger than for other forms of SIS^10,29^. In neither case was PRQ increased relative to non-heat-shocked controls **(Supplementary Fig. S6c,d)**.

It is possible that some of our measurements were statistically underpowered to observe PRQ. It is also possible that for some stressors, PRQ may occur or continue after the 12 h duration of our recordings.

While PRQ is enhanced by EGF signaling and by UV exposure, we did not find statistically significant upregulation of PRQ by a variety of cellular stressors that also cause SIS, with the exception of 9-12 h following Cry5b exposure. Our finding that the propensity and timing for PRQ vary widely after different cellular stressors suggests that the behavioral responses to cellular stressors are not well modeled by our EGF overexpression protocol, but instead may depend on the type of stressor in a complex way.

### Post-response quiescence requires neuropeptide signaling and the sleep active neurons ALA and RIS

We asked whether PRQ, a type of behavioral quiescence, depends on neurons required for other quiescent behaviors. The sleep-active interneurons ALA^12^ and RIS^37^ regulate *C. elegans* behavioral quiescence, and RIS has been shown to be activated during mechanosensory deprivation of lethargus quiescence^38^. The homeodomain transcription factors CEH-14 and CEH-17 are required for proper differentiation of the ALA neuron^39^, and loss of function of either of these genes causes a near total loss of EGF-induced quiescence^9,39^. Another transcription factor, APTF-1, is required for the quiescence-promoting function of RIS^37^. In *aptf*-1 mutants, movement quiescence during lethargus is severely curtailed^37,40^. The RIS neuron plays a role in several other forms of quiescence, including fasting quiescence, satiety quiescence, quiescence associated with early larval developmental diapause^41^, and SIS^12^.

To test whether ALA or RIS play a role in PRQ, we overexpressed EGF in *ceh-14, ceh-17*, and *aptf-1* backgrounds. EGF-induced quiescence was greatly reduced in each of these mutants **(Fig. 4a, Supplementary Fig. S7a)**. These results show that both ALA and RIS regulate EGF-induced quiescence, as previously reported^9,12^.

**Figure 4.**
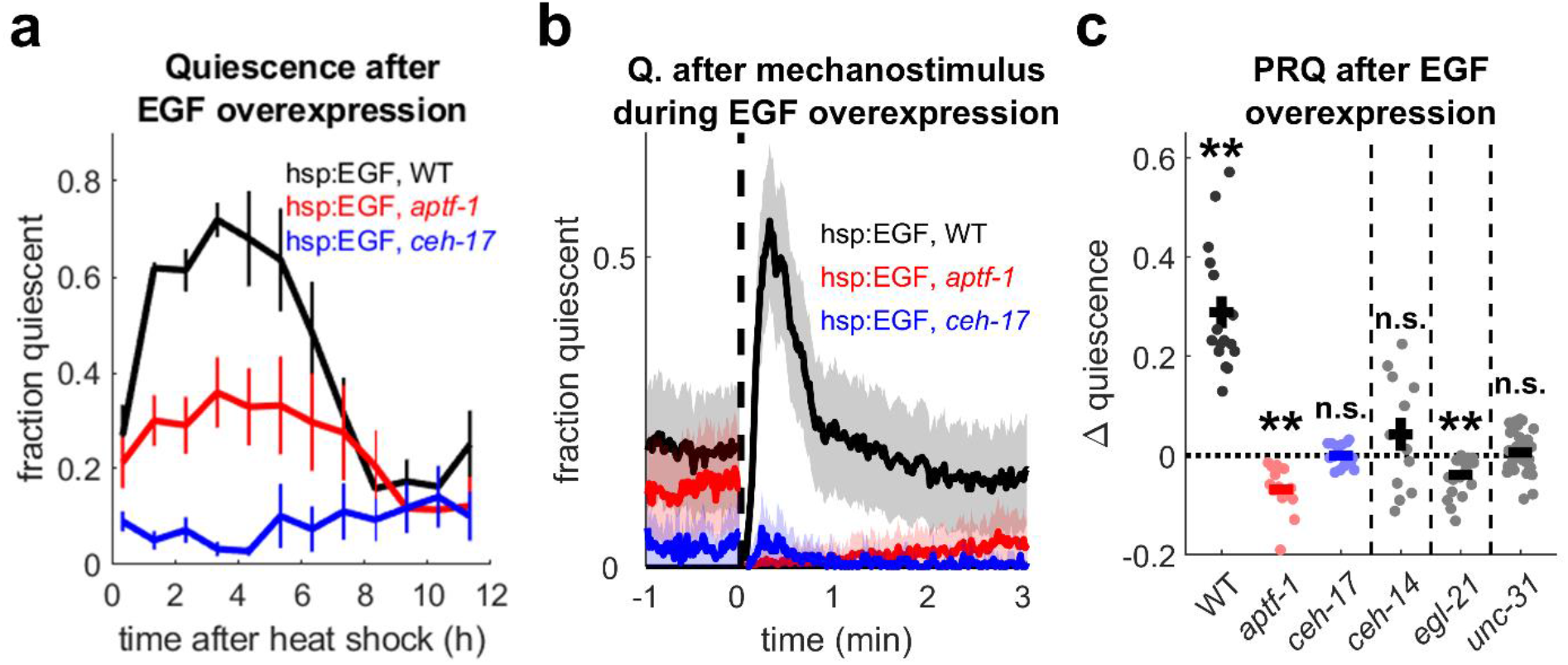
The ALA and RIS neurons and neuropeptide signaling are required for PRQ. **(a)** Fraction of time quiescent of WT (black), *apft-1* (red), and *ceh-17* (blue) mutant animals following EGF overexpression in the absence of additional stimuli (quiescence averaged in 1 h bins, n = 8 animals per genotype, one replicate). Error bars represent ± SEM. **(b)** Fraction quiescent before and after mechanostimulus (vertical dashed line) in WT (black) and *aptf-1* (red) and *ceh-17* (blue) mutant background animals carrying the hsp:EGF array 2-12 h after mild heat shock. n = 16 animals per genotype, two replicates. Shading represents ± SEM. **(c) (left)** Quantification of PRQ in WT and *aptf-1* and *ceh-17* mutants in panel b and **(right)** *ceh-14, egl-21*, and *unc-31* mutants. Δ quiescence was calculated as in Fig. 1a. Dots represent individual animals. n = 12 animals, 2 replicates for *ceh-14*; n = 24 animals, 2 replicates for *egl-21*; and n = 36 animals, 3 replicates for *unc-31*. ** denotes p < 0.01 (two-sided Wilcoxon signed-rank test with Bonferroni correction for three comparisons, left, one sample two-tailed t-test, no correction, right). Error bars represent ± SEM.

In the period 2-12 h after induction of EGF overexpression, neither *ceh-17* nor *aptf-1* mutants showed an increase in quiescence following mechanical stimulus, suggesting that both ALA and RIS are required for PRQ. However, we observed a difference in the behavior of these mutants: *ceh-17* and *ceh-14* mutant animals showed a brief recovery of quiescence to baseline levels within the first minute after the mechanical stimulus, but this was followed by several minutes of reduced quiescence **(Fig. 4b,c, Supplementary Fig. S7b)**. In contrast, *aptf-1* mutants responded to the vibration by briefly moving and then gradually returning to baseline quiescence levels over the course of several minutes. As a result, *aptf-1* mutants showed a significant *decrease* (p = 0.0004) in peak quiescence in the two minutes following the stimulus **(Fig. 4c)**. These results suggest different roles for ALA and RIS in regulating PRQ depth and dynamics.

ALA and RIS regulate sleep and quiescent behavior through the release of sleep-promoting neuropeptides, including those encoded by *flp-13*,^13^ *flp-24*, and *nlp-8* in ALA^14^, and *flp-11* in RIS^40^. A neuropeptide encoded by *nlp-22* is expressed in the RIA interneuron and also promotes quiescence^42^. We therefore asked whether PRQ is also regulated by neuropeptide signaling. EGL-21 is a carboxypeptidase^43^ required for the processing of members of both the FMRF-amide-like (FLP) and neuropeptide-like (NLP) neuropeptide families, and UNC-31 is the *C. elegans* homologue of the calcium-dependent activator protein for secretion (CAPS), which is required for dense core vesicle release^44^, and therefore normal secretion of neuropeptides. To test whether neuropeptide signaling regulates PRQ, we studied animals overexpressing EGF in *egl-21* and in *unc-31* mutant backgrounds.

Mutants of *egl-21* and *unc-31* showed reduced EGF-induced quiescence compared to control animals overexpressing EGF in a wild-type background **(Supplementary Fig. S7a)**. Following substrate vibration, the quiescence of *egl-21* mutants stayed below baseline levels for several minutes, similar to the observation in *aptf-1* mutants. *unc-31* mutants, while also lacking PRQ, quickly returned to baseline quiescence levels **(Supplementary Fig. S7b)**. Because *unc-31* animals move sluggishly^45^ and have reduced touch sensitivity (M. Chalfie, personal communication), we asked if a reduced movement response to mechanical stimuli might explain their lack of PRQ. To address this question, we restricted our comparison of PRQ to episodes in which *unc-31* mutants had a movement response to mechanical stimulus that was similar in magnitude to that of wild-type controls. We found that *unc-31* animals with mechanosensory response activity similar to that of WT animals were still defective in PRQ **(Supplementary Vid. 4, Supplementary Figs. S8, S9)**. These results show that the lack of PRQ in *unc-31* animals is not a product of their reduced movement response to mechanical stimuli.

In addition to expressing FLP-11, RIS also expresses markers of GABAergic neurons^40,46^. We first asked whether GABA signaling is required for SIS by measuring quiescence after UV exposure in *unc-25* mutants. The gene *unc-25* encodes a glutamate decarboxylase required for GABAergic neurotransmission^47^. Overall quiescence following EGF overexpression was similar between *unc-25* and wild-type controls **(Supplementary Fig. S10a)**. We asked whether GABA signaling plays a role in PRQ. *unc-25* mutants showed a decrease in PRQ as measured by change in peak quiescence, and a slight change in shape of the PRQ peak, suggesting that GABA plays a role in regulating the depth and dynamics of PRQ **(Supplementary Fig. S10b,c)**. Indeed, recently published work on the role of RIS as a “stop” neuron show delayed stopping dynamics upon RIS activation in a different GABA transmission mutant^48^. These findings suggest that the regulation of quiescence in response to stress and the regulation of PRQ are controlled by overlapping but distinct mechanisms.

Taken together, these results show that PRQ is regulated by neuropeptide and GABA signaling and by the quiescence-promoting interneurons ALA and RIS.

### Post response quiescence does not fulfill several expectations and criteria for sleep or sleep homeostasis

Sleep can be defined behaviorally as a quiescent state with three additional characteristics: (1) reduced responsiveness to stimuli, (2) rapid reversibility^49^, and (3) homeostatic regulation^50^. Animals in SIS have been shown to be less responsive to stimuli and SIS has been shown to be rapidly reversible^10,29^. However, despite being considered a sleep state, to our knowledge SIS has not been shown to be under homeostatic regulation. Because PRQ occurs following a disruption of SIS quiescence, and requires some of the same neurons and genes as two previously described *C. elegans* sleep states (SIS as well as lethargus, or developmentally-timed sleep^16^) we asked whether PRQ might also be a form of sleep homeostasis. We performed several experiments to test this idea.

Homeostatic regulation of lethargus has been characterized previously^31,51,52^. First we considered potential similarities between homeostatic regulation of lethargus and the PRQ we observe during SIS. All sleep homeostasis manifests as an increase in sleep or sleep drive in compensation for deprivation of sleep. *C. elegans* lethargus consists of alternating bouts of activity and quiescence, and long-term (30 min – 1 h) interruption of these quiescent bouts causes an extended period of increased quiescence lasting the remainder of the duration of lethargus^31,51^, whereas brief stimuli cause a short term increase in quiescence on a time scale similar to PRQ^52,53^. Thus we asked whether PRQ during SIS is similar to short-term homeostatic rebound in lethargus.

Short-term homeostasis is associated with a positive correlation between the durations of quiescent bouts and the preceding active bout, even in the absence of experimental stimuli, a phenomenon that has been named microhomeostasis^52,53^. Genes required for microhomeostasis are also required for short-term homeostasis^52^. First, we reproduced this result, finding a significant (p = 0.01) positive correlation between the durations of quiescent and active bouts during lethargus **(Fig. 5a)**. This demonstrated our ability to detect microhomeostasis in our experimental setup.

**Figure 5.**
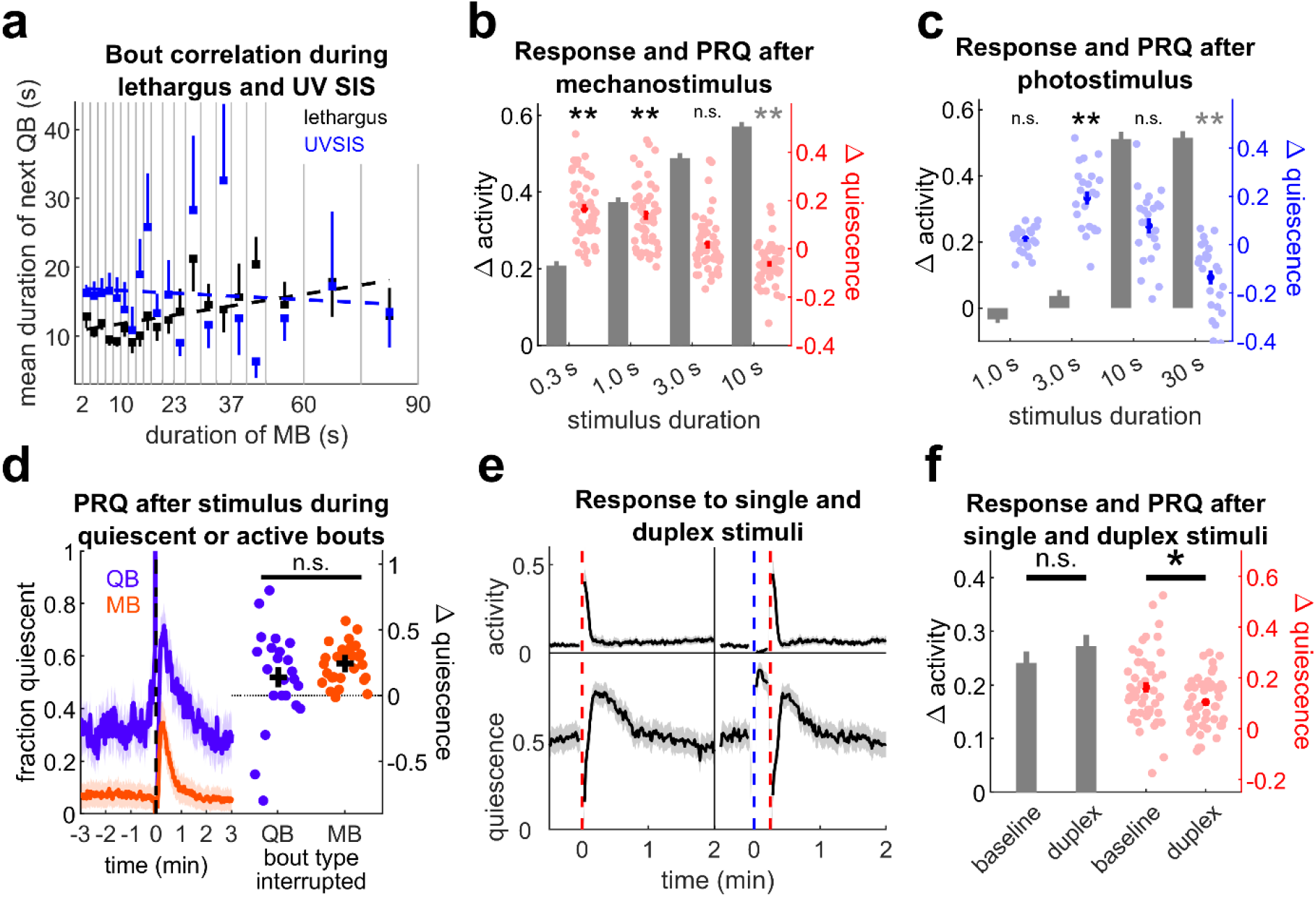
Post-response quiescence lacks some characteristics of sleep homeostasis. **(a)** Correlation of the durations of quiescent bouts (QBs) with the preceding movement bout (MB) duration in lethargus (black) and UV SIS (blue). Squares represent average QB duration following MBs in each MB duration bin, bars represent SEM. Bout duration was significantly correlated for lethargus (slope = 0.089, p = 0.01, R^2^ = 0.32), but not UV SIS (slope = −0.028, p = 0.69, R^2^ = 0.0090). n = 17 animals, three replicates for lethargus, n = 12 animals, two replicates for UV SIS. **(b)** Mechanosensory response (gray bars) and PRQ (red) of WT background animals 2-12 h after EGF overexpression. **\*\*** denotes p < 0.01, two-sided Wilcoxon signed rank test with Bonferroni correction for four comparisons, note that 10 s stimulus animals were *less quiescent* after stimulus. n = 48 animals, four replicates. **(c)** Photosensory response (gray bars) and PRQ (blue) of WT background animals 2-12 h after EGF overexpression. **\*\*** denotes p < 0.01 (two-sided Wilcoxon signed-rank test with Bonferroni correction for four comparisons, note 30 s stimulus animals were *less quiescent* after stimulus). n = 23, two replicates for 1, 3, and 10 s, n = 24, three replicates, for 30 s stimuli) **(d) (left)** Average quiescence of WT background animals 2-12 h after EGF overexpression after mechanostimulus (dashed line) when the stimulus interrupted a quiescent bout (blue) or an active bout (red). n = 28 animals, four replicates. These data also appear in Fig. 2. **(right)** Comparison of PRQ from the left. Data from 4-2 min before the stimulus were used for baseline quiescence. Dots represent single animal averages for stimuli that interrupted quiescent or active bouts. PRQ was not significantly different (p = 0.17, two-tailed, two sample t-test). **(e)** Average normalized activity (top) and quiescence (bottom) following a single 1 s mechanostimulus (left) and duplex (15 s separating a 3 s blue light stimulus and 1 s mechanostimulus) (right) stimulus in WT background animals 2-12 h after EGF overexpression. Red and blue dashed lines represent mechanostimuli and blue light stimuli, respectively. n = 47 animals, two replicates. These responses are quantified in panel f. **(f)** Mechanosensory response (gray bars, left) of WT background animals 2-12 h after EGF overexpression after single and duplex stimuli, and PRQ (red, right) of the same animals after single and duplex stimuli. * denotes p < 0.05 (two sample t-test). Δ quiescence was calculated as in Fig. 1a. Dots represent individual animal averages and error bars and shading represent ± SEM.

We next examined the bout architecture of UV SIS to determine if, like lethargus, it also exhibits microhomeostasis. During UV SIS, quiescent bout durations were not significantly correlated with the preceding movement bout duration **(Fig. 5a)**. There was a weak positive correlation (correlation coefficient 0.39) between quiescent and movement bouts in EGF-induced quiescence, but this correlation did not reach statistical significance (p = 0.089 **(Supplementary Fig. S11)**. These results suggest that PRQ, while resembling short-term lethargus homeostasis, is not associated with microhomeostasis, and therefore may be mechanistically and teleologically distinct.

Next we asked what characteristics PRQ might have if it were a form of sleep homeostasis. In other animals, sleep duration and/or depth increases with the duration of prior wakefulness^54^. If PRQ is a form of sleep homeostasis, then increasing the amount of quiescence interruption should increase the amount of PRQ. We performed several experiments to test this prediction. First, we induced EGF overexpression in WT-background animals and applied aversive mechanosensory or blue light stimuli every 15 minutes. Unlike in previous experiments, we varied the duration of these stimuli to elicit different magnitudes of response, ranging from little or no increased activity following the weakest stimuli, to larger increases in activity after longer stimuli. However, increased response was associated with a *decrease* rather than an increase in quiescence, suggesting that PRQ does not increase with greater interruption of SIS **(Fig. 5b,c, Supplementary Fig. S12)**. However, we note that while PRQ is likely distinct from the short-term homeostasis described by Nagy *et al*.^52^, they did report that stronger stimuli did not cause a short term increase in quiescence, but rather a long term increase during the entire inter-stimulus interval (15 min).

Next, we examined responses to 1 s substrate vibration and compared the increase in quiescence in animals in which the stimulus occurred during a quiescent bout to that in animals in which the stimulus occurred during an active bout. In a study of long-term deprivation of developmental lethargus, Driver *et al*. reported increased mortality in sleep deprivation susceptible *daf-16* mutants stimulated during quiescent bouts, but not in yoked controls that were stimulated simultaneously regardless of whether they were quiescent^51^. Although similar experiments measuring amount of homeostatic compensation have not been reported, we reasoned that if PRQ were sleep homeostasis, then PRQ magnitude would be larger in cases where the stimulus disrupted a bout of quiescence compared to cases in which the stimulus occurred during a movement bout. We did not observe a significant difference in PRQ between these groups (p = 0.17) **(Fig. 5d)**. These results suggest that PRQ is not a homeostatic compensation for interruption of quiescence. While we sought to compare PRQ and short-term homeostasis directly, we were unable to replicate the increase in quiescence in L4 lethargus animals following mechanostimulus reported by Nagy *et al*.^52^ **(Supplementary Fig. S13)**.

Lastly, we reasoned that if PRQ were a homeostatic compensation for interruption of SIS, it must itself be sleep and therefore feature rapid reversibility, decreased response to stimuli, and homeostatic regulation of itself^49,50^. To determine whether PRQ fulfills these criteria, we first induced PRQ using either a 3 s blue light pulse. We then applied a 1 s mechanostimulus 15 s later, near the peak of PRQ. The animals moved in response to the second stimulus, indicating that PRQ is reversible. To determine whether responsiveness is decreased in PRQ, we compared the stimulus evoked activity during PRQ to the stimulus evoked activity following single mechanostimuli not preceded by a PRQ-inducing stimulus in the same animals. We observed a small, non-significant increase in response activity after the second stimulus compared to the single stimulus, indicating that animals in PRQ are at least as responsive to stimuli as the same animals outside of PRQ. This result suggests the PRQ does not represent an increase in sleep, which is a low arousability state. To determine whether PRQ is under homeostatic regulation, we looked for rebound quiescence following the stimulus duplex. While we observed a second peak in quiescence following the stimulus duplex, the PRQ following the stimulus duplex was significantly less than PRQ following a single mechanostimulus (p = 0.022), despite higher levels of quiescence at the time of the second stimulus, suggesting that PRQ itself is not homeostatically regulated **(Fig. 5e,f)**.

These results show that PRQ is not associated with the microhomeostatic bout correlations seen in similar homeostasis of another form of *C. elegans* sleep, that PRQ is not correlated with the amount of quiescence deprivation, and that PRQ lacks some of the criteria for sleep, reduced arousability and probably homeostasis. Therefore, PRQ does not fulfill many expectations for an SIS homeostasis mechanism, and homeostatic regulation of SIS remains to be demonstrated.

### During PRQ, lateral head movements are suppressed during locomotor pauses

Quiescent states often involve the coordinated suppression of multiple behaviors. Different aspects of SIS quiescence – locomotion, head movement, feeding, and defecation – are regulated by different combinations of neuropeptides^14^. On the other hand, active behaviors can also involve inhibition of movements, such as in the suppression of lateral head movements during the escape response^28,55^. We asked how different types of behavioral quiescence may be coordinated during PRQ.

Our image subtraction-based analysis in the WorMotel could not distinguish between head movement and whole-body locomotion^56^ or detect pharyngeal pumping, an indicator of feeding^57^. To assay these more subtle behaviors we recorded videos with higher spatial and temporal resolution of the mechanosensory responses of individual animals on agar pads before and after EGF induction. We then manually scored lateral head movements (movements independent of overall body movement and including movements of only the nose), forward and reverse locomotion (mid-body movement relative to the substrate), and pharyngeal pumping.

We first sought to verify that EGF-induced quiescence and PRQ could be recapitulated in this agar pad preparation. Consistent with previous reports^9,14^, EGF overexpression decreased overall locomotion, head movement, and pumping rate **(Supplementary Fig. S14)**. In our WorMotel analysis, quiescence is defined as lack of movement throughout the body, and therefore implies simultaneous quiescence in both locomotion and head movement. To compare our WorMotel data to our higher resolution agar pad data, we defined simultaneous locomotion and head movement quiescence as “body quiescence”. We found a significant increase in body quiescence after the stimulus (p = 0.015) with a time dependence comparable to our WorMotel data for PRQ **(Fig. 6a,b)**. These results confirm that worms exhibit PRQ in our high-resolution assay.

**Figure 6.**
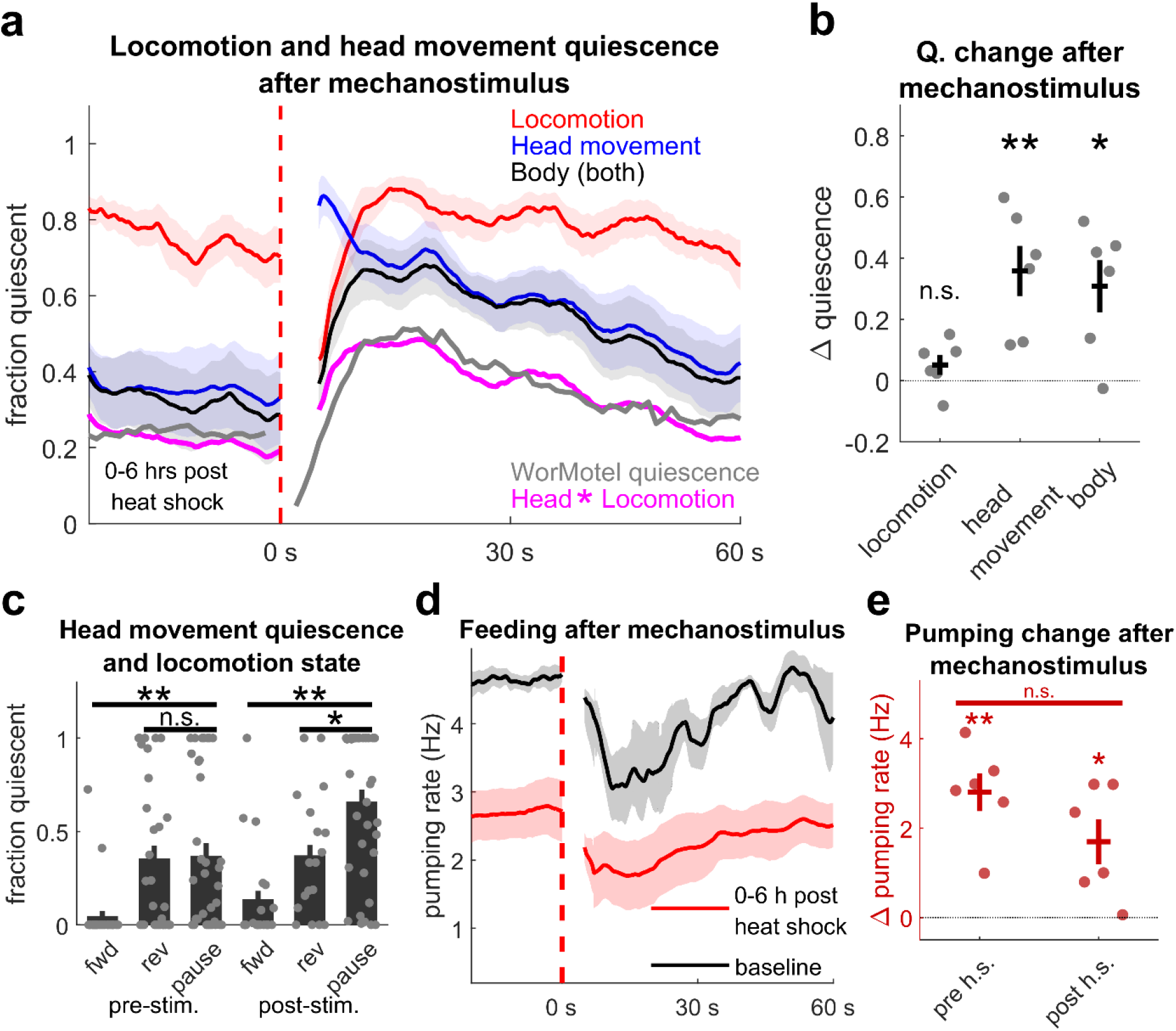
Locomotion, head movement, and feeding during PRQ. **(a)** Traces showing fraction of time animals were locomotion, head movement, and body movement quiescent (both locomotion and head movement quiescent) after mechanostimulus following EGF overexpression. Traces represent the mean of six animals, four recordings each, 0-6 h after heat shock, smoothed with a 5 s window. The magenta trace is the product of the locomotion and head movement traces, and the gray trace is quiescence following mechanostimulus measured in the WorMotel 1-6 hours after EGF overexpression (same data shown in Fig. 2b). The dashed red line indicates the stimulus. Data from the same animals are used throughout Fig. 6 and Supplementary Fig. 14. **(b)** Average change in peak locomotion, head movement, and body quiescence (black) and average change in minimum pumping rate (red) from the 25 s period before the stimulus to the 25 s period after the stimulus starting at 5 s post stimulus. Dots represent individual animal averages. * denotes p < 0.05, ** denotes p < 0.01, two-tailed, one sample t-test. Data are the same as panels a-c, 1-6 h post heat shock. **(c)** Fraction of time the animals were head movement quiescent while moving forward, reversing, or pausing before (left) and after (right) mechanostimulus. Dots represent single recording averages. * denotes P < 0.05, ** denotes p < 0.01, two-sided Wilcoxon rank sum test. **(d)** Traces showing mean pumping rate before heat shock (black) and 1-6 hours after heat shock (red) after mechanostimulus (dashed line) in hsp:EGF animals. Traces are smoothed with a 5 second averaging window and represent 6 animals, one recording each prior to heat shock, six recordings each 1-6 hours after heat shock. **(e)** Average change in minimum pumping rate from the 25 s period before the stimulus to the 25 s period after the stimulus starting at 5 s post stimulus, before and after heat shock. Dots represent individual animal averages. * denotes p < 0.05, ** denotes p < 0.01. Two tailed one sample or paired sample t-tests were used. Error bars and shading represent ± SEM.

Next, we examined the two components of body quiescence, head movement and locomotion quiescence. To our surprise, we observed no significant increase in locomotion quiescence after vibration in EGF induction animals, suggesting that locomotion quiescence does not increase during PRQ. In contrast, we found a significant increase in head movement quiescence after mechanostimulus (p = 0.007) **(Fig. 6b)**, indicating that head movement, rather than locomotion, is suppressed during PRQ.

If the timing of head movement quiescence and locomotion quiescence bouts was statistically independent, we would expect the product of the fractions of head movement and locomotion quiescence to be equal the fraction of body quiescence. In fact, we found that body quiescence fraction was greater than the product of the head movement quiescence and locomotion quiescence fractions. Indeed, beginning about 10 s after the mechanostimulus, head movement and body movement quiescence are at similar levels **(Fig. 6a)**. This observation suggests that during the timeframe of PRQ, head movement quiescence specifically occurs while the animal is undergoing locomotion quiescence. Consistent with this observation, comparison of the rate of head movement quiescence during forward, reverse, and pause locomotion states, before and after mechanostimulus, shows that head movement quiescence was greatest when the animal was pausing after mechanostimulus **(Fig. 6c)**.

Using our high-resolution assay, we also asked to what extent feeding is suppressed during PRQ. We found that pumping rate decreased after the mechanostimulus, consistent with findings that mechanosensory stimuli inhibit pumping^21,58^. However, the magnitude of decrease in pumping rate following mechanostimulus was similar before and after EGF induction **(Fig. 6d,e)**. This finding suggests that feeding behavior does not display an EGF-inducible quiescence following mechanostimulus similar to the suppression of body movement during PRQ.

These results suggest that during PRQ, locomotion and head movement quiescence are coordinated such that head movement quiescence occurs during locomotor pauses.

## Discussion

In this study we have described PRQ, a form of quiescence that follows *C. elegans* responses to mild, aversive stimuli. This behavior is distinct from other forms of quiescence previously described in *C. elegans*. For example, animals touched in the anterior part of the body will sometimes exhibit an immediate and brief pause in forward movement^59^. Compared to these pauses, PRQ is much longer in duration and occurs with a delay relative to the stimulus. As another example, animals confined to small spaces engage in long-term quiescent behavior^60^. PRQ occurs more quickly and is much shorter in duration than this confinement induced quiescence.

Several lines of evidence suggest that PRQ is related to SIS. PRQ is potentiated following UV SIS and following overexpression of EGF, which mediates the response to heat shock and probably other stressors^10^, and PRQ and SIS share a dependence on the ALA and RIS quiescence-promoting interneurons and neuropeptide signaling. However, PRQ may not be a direct consequence of SIS, as some forms of stress induce SIS but not PRQ. Furthermore, the time course of PRQ following stress or EGF overexpression is distinct from that of SIS, with PRQ reaching a maximum after that of SIS and continuing many hours after cessation of the SIS response. While we cannot rule out a role of low level leaky EGF expression in the persistence of PRQ long after EGF overexpression, this would not explain the persistence of PRQ for many hours following UV exposure.

We asked what might be the adaptive significance of the PRQ behavior. Here we consider three ideas: (1) that PRQ reflects a homeostatic mechanism in SIS (2) that PRQ results from energy conservation, and (3) that PRQ is a form of defensive freezing.

Given the association between PRQ and SIS, we first considered the possibility that PRQ represents a homeostatic mechanism in SIS, or rebound SIS. However, during PRQ, animals do not exhibit reduced arousability and we did not find evidence for PRQ being under homeostatic regulation. Therefore we did not observe the properties expected for PRQ to be a sleep state.

A second possible interpretation of PRQ is in terms of energy conservation. *C. elegans* locomotion requires energy expenditure^61^, and cessation of feeding and activation of DNA repair pathways may contribute to reduced energy stores during UV SIS^62^. Indeed, fat stores decrease during lethargus and ATP levels decrease during lethargus and over the course of UV SIS^62^. PRQ may allow the animal to compensate for energy depletion associated with an active response with cessation of locomotion and feeding. However, we consider this energy-based argument unlikely since longer stimuli elicit more activity than brief stimuli without causing PRQ.

A third possibility is that PRQ represents a form of defensive freezing, analogous to the freezing phase in the classic mammalian escape response^4,63^. A cessation of movement can help an animal that has detected a predator to evade detection by the predator^4,64^. Given the limited visibility in *C. elegans*’ natural environment of decaying organic material, it is likely that its predators rely on primarily olfactory, mechanosensory, and potentially electrosensory^65^ cues rather than visual ones for prey seeking and identification. *C. elegans’* natural predators likely include nematophagous arthropods, fungi, and other nematodes^28,66,67^. PRQ may allow *C. elegans* to avoid detection after retreating from contact with a predator by minimizing mechanosensory stimulation, electromyogenic cues, and potentially olfactory cues, since touch suppresses defecation^68^. Reduced feeding during PRQ (and after touch generally) may help minimize the ingestion of harmful fungal or bacterial spores, and the suppression of head movement during PRQ suggests it may be related to the tyramine-dependent suppression of head movement during *C. elegans* escape behavior^28,55^.

Additional lines of evidence support an association between PRQ and defensive behaviors. We found that RIS function is required for PRQ, and *unc-25* (GABA synthesis) mutants show a partial deficit in PRQ. GABAergic function in the RIS neuron has been found to regulate *C. elegans* avoidance of kairomones released by the predatory nematode *Pristionchus pacificus*^69^, supporting a connection between PRQ and defensive avoidance behavior.

It is unclear why defensive freezing would be upregulated after *cellular* stress. In fact, we observe PRQ more reliably following EGF overexpression than after any stressor tested, suggesting that EGF, not cellular stress, controls PRQ in *C. elegans*. In other organisms, EGF signaling plays a role in psychological, not just physiological stress. In mice, the EGF-family receptor neuregulin-1 has been shown to modulate anxiety-like behaviors^70,71^; for example, overexpression of the neuregulin-1 receptor increases baseline startle response^72^. These findings suggest that EGF family signaling may play conserved roles in regulating defensive and fear/anxiety behaviors.

In summary, PRQ represents a novel feature of *C. elegans* escape behavior during a stress state. Further study of its circuit and genetic bases will enrich our understanding of the mechanisms modulating animal responses to aversive stimuli.

## Methods

### C. elegans strains & maintenance

We cultured *C. elegans* on OP50 *E. coli* food bacteria on standard NGM agar plates^32^ at 20°C, and experiments were performed at ambient lab temperature, which ranged over 18°C – 24°C. Unless otherwise specified, we performed experiments using young adult hermaphrodites staged by picking late L4 animals 5 – 7 hours before the experiment. The exceptions to this are the lethargus experiments, in which we used mid to late L4 larvae picked just before the experiment.

Strains used in this study and the phenotypes used for identifying mutant progeny are listed in Table 1. We used the Bristol N2 strain as wild type. For EGF overexpression experiments in a wild-type background, we crossed PS5009 *pha-1(e2132ts)* III.; *him-5(e1490)* V. syEx723[*hsp16-41*::*lin-3c*; *myo-2*:GFP; *pha-1*(+)]^9^ with N2 to make YX256. For EGF overexpression experiments in other mutant backgrounds, we crossed mutant strains with YX256 males to generate strains carrying *syEx723* in the mutant background. We used the phenotype indicated in the table below to identify mutant progeny. All mutants used to generate crosses are available from the CGC.

**Table 1.** Strains used in this study and phenotypes used to identify mutants.

| Strain | Genotype | Phenotype | Cross |
| --- | --- | --- | --- |
| Bristol N2 | WT | - | - |
| YX256 | *syEx723* [*Phsp16.41::lin-3c*; *Pmyo-2::GFP*; *pha-1(+)*] | - | PS5009 x N2 |
| YX271 | *unc-31(e928)* IV; syEx723 | Uncoordinated | YX256 x DA509 |
| YX276 | *aptf-1(tm3287)* II; syEx723 | Reduced lethargus quiescence | YX256 x HBR232 |
| YX277 | *ceh-17(np1)* I; syEx723 | Reduced UV SIS quiescence | YX256 x IB16 |
| YX278 | *unc-25(e156)* III; syEx723 | Uncoordinated | YX256 x CB156 |
| YX280 | *unc-31(e169)* IV; syEx723 | Uncoordinated | YX256 x CB169 |
| YX281 | *ceh-14(ch3)* X; syEx723 | Reduced UV SIS quiescence | YX256 x TB528 |
| YX282 | *mec-4(u253)* X; syEx723 | Mechanosensory defect | YX256 x TU253 |

### Heat shock, UV, and other stressors

#### UV

To expose worms in WorMotels to UV light, we used a Spectrolinker XL-1000 UV Crosslinker (Spectronics Corporation, Westbury, NY) with UV-C (254 nm) fluorescent tubes. The WorMotel was placed uncovered on a flat black background in the bottom middle of the crosslinker, and the crosslinker was run at the indicated energy dose from a cold start (after having been off for at least several hours).

#### Heat

For most experiments, we used a thermal immersion circulator to maintain a water bath at the specified temperature. For some experiments, we used a hot plate and added hot or cold water to maintain the temperature. We fully immersed a Parafilm-sealed petri dish containing the WorMotel for the specified time, then placed the petri dish on the lab bench for about 5 min to return to ambient temperature before proceeding.

#### Cold

We chilled a Parafilm-sealed petri dish containing the WorMotel and young adult worms in a 4°C refrigerator or 4°C thermal circulator for 24 hours.

#### Salt

We picked WT worms from a WorMotel into NGMB (NGM buffer) containing an additional 500 mM NaCl for 15 minutes and then picked them back onto the WorMotel using an eyebrow pick. Controls for this experiment were picked into regular NGMB without extra salt. NGMB has the same composition as Nematode Growth Medium (NGM)^73^ but without agar, peptone, and cholesterol.

#### Ethanol

We picked WT worms from a WorMotel into a 5% (v/v) solution of ethanol in NGMB for 15 minutes using an eyebrow pick. Controls for this experiment were picked into NGMB without ethanol.

#### Cry5b

We transferred WT worms from the WorMotel onto a standard NGM plate seeded with Cry5b-expressing bacteria^10,74^, left the animals on the Cry5b-expressing bacteria for 15 minutes, and then transferred them back into individual wells of the WorMotel. Controls were picked onto a plate seeded with bacteria expressing the vector backbone of the Cry5b-expressing plasmids.

### Mechanosensory stimulation

For mechanosensory assays performed while imaging using dark field illumination, we coupled the WorMotel to an audio loudspeaker (PLMRW10 10-Inch Marine Subwoofer, Pyle Audio Inc., Brooklyn, NY). Acrylic mounting plates with screws fixed the WorMotel tightly to the inside of a 10 cm petri dish, which itself was mounted on the center of the speaker cone using an acrylic ring and screws. The field was illuminated with a ring of red LEDs situated around the Petri dish.

For mechanostimulus assays performed with bright field imaging for scoring pharyngeal pumps, we coupled a 3 mm thick agar pad contained in a custom acrylic pad holder to a BRS40 4-inch audio loudspeaker (BOSS Audio Systems, Oxnard, CA) using a 50 mL polystyrene serological pipette glued to the middle of the speaker cone and projecting horizontally. A notch was cut into the end of the pipette tip to fit snugly onto a tab on the pad holder, transferring vibrations from the speaker to the worm substrate.

For all mechanosensory assays, we used a custom MATLAB (MathWorks, Natick, MA) script to output audio signals though either a NIDAQ PCI-6281 (National Instruments, Austin, TX) or the computer’s audio jack to a KAC-M1804 Amplifier (Kenwood, Long Beach, CA), which powered the loudspeaker. Audio signals from the NIDAQ were 1 kHz at a 1 V 0 to peak amplitude except where noted. The beginning and end of the envelope of the audio waveform were made smooth by initiating and terminating using two halves of a 0.1 second Hann window. Waveform duration varied by experiment as specified.

### Blue light stimulation

The blue light setup is similar to one previously described^30^. We used a high power blue LED (Luminus PT-121, Sunnyvale, CA, center wavelength 461 nm) driven at 20 A using a DC power supply to illuminate the worms with 0.36 mW / mm^2^ blue light. Except when combining mechanosensory and photosensory stimuli in the same experiment, we placed the WorMotel and LED inside a box made of mirrored acrylic interior walls to improve illumination uniformity. A custom MATLAB script controlled illumination timing through a NIDAQ USB-6001 (National Instruments, Austin, TX) data input/output device and relay (6325AXXMDS-DC3, Schneider Electric, France).

### WorMotel fabrication and preparation

We fabricated WorMotels as described previously^30^. Briefly, we poured PDMS (Dow Corning Sylgard 184) into 3D-printed acrylic or polycarbonate molds, cured them overnight at 50 °C, and then removed the PDMS WorMotel devices. We boiled WorMotels in DI water to sanitize them, baked them in a 50°C oven to dry, and treated them with oxygen plasma for 10 seconds in a plasma oven consisting of a Plasma Preen II 973 controller (Plasmatic Systems, Inc.) connected to a microwave oven (Amana RCS10TS). Plasma treatment makes the PDMS device hydrophilic, rendering it easier to fill and the agar surfaces flatter.

We used a pipette to fill the wells with either media containing 3 g/L low melt agarose (Research Products International, Mount Prospect, IL), 5 mg / L cholesterol, 2.5 mg / L bacto-peptone, 1 mM CaCl_2_, 1 mM MgSO4, 20 mM KH_2_PO_4_, and 5mM K_2_HPO_4_) (variable duration blue light and vibration experiments) or standard NGM agar^32^ (all other experiments). To reduce the rate of worms escaping from their wells, we filled the moats surrounding the wells with an aversive solution of 100 mM CuSO_4_^30^.

We seeded agarose-filled WorMotels with 5 μL of an overnight *E. coli* OP50 culture resuspended in 3g / L NaCl and allowed the bacteria to dry, re-wetting faster drying wells with DI water to ensure even drying overall, and either used the WorMotels immediately or stored them in a Parafilm-sealed dish with hydrated water crystals (AGSAP PAM, small particle size, M2 Polymer, 1.5 g in 500 mL water) at 4°C for up to two weeks. We seeded agar-filled WorMotels with 2 μL of an overnight culture of OP50 in LB, and aspirated excess LB before allowing the WorMotels to dry open in a biosafety cabinet for 8 minutes. We incubated these WorMotels in a Parafilmed dish with hydrated water crystals (150:1 water:crystals) for three days at 20°C, allowing a bacterial lawn to grow. We either used the WorMotels immediately or stored them at 4°C for up to two weeks until needed.

After each experiment, we emptied the wells of agar / agarose, washed the WorMotels with hot water and detergent (Alconox), rinsed in DI water, dried, and stored them for reuse.

### Imaging and image processing in the WorMotel

For imaging in the WorMotel, we used a 10.7 MP CMOS camera (DMK 24UJ003, The Imaging Source, Charlotte, NC) and 12.5 mm lens (HF12.5SA-1, Fujifilm Corp., Tokyo, Japan) controlled by MATLAB to acquire 8 bit, 10 MP grayscale images of worms in WorMotel wells at 1 frame per second. Red (wavelength 628 nm) LED strips (Oznium, Pagosa Spring, CO) provided dark field illumination. For most experiments, we imaged 24 wells (a 4 x 6 array) at a resolution of approximately 7.4 μm per pixel. The exceptions to this were data used in bout correlation analysis, where we used the same camera with a 50 mm lens (Fujinon HF50SA-1, Fujifilm Corp.) to image six wells (a 2 x 3 array) at approximately 3.6 μm per pixel for L4 lethargus experiments or 12 wells (a 3 x 4 array) at approximately 5.0 μm per pixel for young adult SIS and EGF-induced quiescence experiments.

We used custom MATLAB scripts for image processing. We smoothed consecutive frames with a Gaussian filter (σ = 1.5 pixels) and then subtracted them. We binarized the absolute value of the resulting difference image using a grayscale threshold value of 5. The number of non-zero pixels in each well ROI in the binarized difference image was summed to determine the activity of each worm. A worm was considered quiescent if no activity was detected in its well region of interest for one frame, and active otherwise. The only exception was for bout correlation analysis, where we used a 2 second floor on quiescent bouts in keeping with the literature^52,53^. To calculate normalized activity for each worm, we divided the activity value of each frame pair by the average activity from the 50 most active frames after excluding the top 0.5%. Stimulus frames and stimulus-adjacent frames were excluded from analysis, as were wells containing no worms, more than one worm, or in which the worm escaped the well or burrowed into the agar substrate during recording. Four worms that remained immobile and non-responsive for the entire recording period following cold shock were also censored.

### Quantification of stimulus response (Δ activity)

Responsiveness was determined by first calculating the average activity of each animal over all stimuli in the relevant time period, then taking the average activity for five seconds following the stimulus (excluding the stimulus frame and one frame after to avoid the artifact from the mechanostimulus), and subtracting the baseline average activity in the minute prior to the stimulus. The only exception was for results shown in Fig. 5f, where the average activity in the 5 s before the stimulus was used as the baseline because of the blue light stimulus preceding the mechanostimulus. We excluded the stimulus frame and the frame before it to again avoid the artifact from the mechanostimulus. When examining how SIS and EGF overexpression affect arousability, responses were normalized to those of controls (untreated in UV SIS experiments or WT in EGF overexpression experiments).

For quantifying the responses of *unc-31* animals and their controls, activity was first calculated by subtraction of non-consecutive frames separated by five seconds, and activity resulting from movement during the stimulus minus average activity in the 20 s prior to the stimulus was used as a measure of the response **(Supplementary Fig. S8,9)**.

### Quantification of SIS and EGF-induced quiescence

To quantify the overall quiescence time course during SIS or after EGF overexpression, we calculated the fraction of time each animal was quiescent during the specified time bin. The y-axes on plots showing this quantity are labeled “fraction quiescent”, the same term used in plots showing PRQ traces using shorter time bins (see below). For experiments with stimuli, we excluded data recorded in the 12 min following a stimulus from this calculation and labeled the y-axis on resulting plots as “baseline fraction quiescent”.

### Quantification of PRQ (Δ quiescence)

We quantified PRQ via the change in quiescence (Δ quiescence) following the stimulus relative to the quiescence before the stimulus. This parameter was calculated by first taking the average quiescence of each animal before and after stimuli in the time period of interest, smoothing this trace with a 10 second averaging filter, and then subtracting the highest quiescence level in any 10 second period in the two minutes before the stimulus from the highest quiescence level in any 10 second period in the two minutes after the stimulus. The only exception was for comparison of quiescent bout-interrupted PRQ to active bout-interrupted PRQ **(Fig. 5d)**, where the baseline period was shifted to two minutes earlier to avoid the increase or decrease in quiescence leading up to the stimulus in the two groups.

### Simulated quiescence traces due to bout synchronization

The traces appearing in Supplementary Figs. S4a and S4b were generated from the PRQ data shown in Fig. 1 and Fig. 2, respectively (see also Supplementary Discussion 2).

To align data by active bouts (top traces, both panels), we examined each animal’s baseline binary activity / quiescence data starting 3 min before each mechanostimulus and stepping backward in time by 1 frame (1 s) increments until finding a quiescent-to-active transition, i.e. the beginning of an active bout. We pooled data aligned by these transitions to create the top traces. For the middle traces, we followed a similar procedure as for the top traces. First, we aligned the baseline data by the beginning of quiescent bouts instead of active bouts. Then we offset each individual worm’s data at the point of alignment by inserting an active bout of the same duration as the response activity bout (the activity bout immediately following the next mechanostimulus in the same animal). See Supplementary Fig. S4c for a schematic of the insertion. We created the bottom traces by aligning the data by the beginning of quiescent bouts without an offset. We excluded data when we did not find the type of bout transition being sought within 12 min before the mechanostimulus, or, for offset traces, if the stimulus response continued until the next stimulus or to the end of the video (14-16% of stimuli for EGF overexpression and 1.2-1.5% of stimuli for UV exposure).

### Imaging and scoring locomotion, head movement, and feeding

To image pharyngeal pumping, we manually tracked unconstrained worms on an agar pad containing OP50 bacteria using an M156 FC stereo microscope (Leica, Wetzlar, Germany) with a 1.0X Plan Apo objective and white, bright field illumination. We used a 5 MP CMOS camera (DMK33GP031, The Imaging Source) to image at 50 fps with a resolution of approximately 700 pixels / mm and a field of view of approximately 0.91 mm x 0.68 mm.

We used custom MATLAB graphical user interfaces to replay images at approximately 15 fps and score individual pumps, the locomotion state (forward, pause, or backward relative to the substrate), and the head movement state (active or quiescent, determined by the presence or absence of lateral movements of the head or nose independent of the rest of the body) of the worm from 30 s before the stimulus to 60 s afterward (see Supplementary Vid. 5). We censored frames where the behavior could not be scored with confidence. The scorer was blind to which group the worm belonged to (pre- or 1-6 h post-EGF overexpression).

### Lifespan measurements

For lifespan measurements, WorMotels were filled with NGM agar as described^30^, except the agar contained 200 μM fluoro-2’-deoxyuridine (floxuridine) to inhibit reproduction, and the WorMotels were seeded with 5 μL of a concentrated suspension of OP50 *E. coli*, which was allowed to partially dry to eliminate liquid on the agar surface. We picked late L4 hermaphrodites into the WorMotel wells, and then treated the UV group as described above. We scored worm survival manually on the days noted. A worm was considered dead if it was not moving or feeding spontaneously and failed to respond to a prod with a platinum wire pick.

### Plotting and statistics

In all scatter plots except those showing movement / quiescent bout correlations, individual dots represent the average behavior of a single animal over the recording period or in the time period specified on the axes and / or in the caption. These single animal averages were used for statistical testing and for calculating the overall average. In plots showing traces, we first computed individual animal traces by averaging the binary score (active or quiescent) of that animal across stimuli in the time period of interest. We then plotted traces representing the average of these traces. This quantity was labeled “fraction quiescent”. Individual animal traces were used to compute the SEM, shown as a shaded area.

For bout correlation plots, we first organized activity / quiescence data into pairs of movement bouts and subsequent quiescent bouts. Then we grouped these pairs into bins according the duration of the movement bout and plotted the mean and standard error of the duration of the subsequent quiescent bouts in each bin.

All statistical tests were performed in MATLAB. Statistical tests and p-values used are listed in the figure captions, as are the number of replicates, which were separate runs of the experiment with different animals on a different day. The Anderson-Darling test was used to test for normality. If any dataset in a set of related experiments was found to be non-normal, we used nonparametric statistical tests for comparisons.

## Additional Information

## Acknowledgements

We thank Raffi Aroian for Cry5b, Paul Sternberg for strain PS5009, and Martin Chalfie and Matthew Kayser for helpful discussions. Some strains were provided by the CGC, which is funded by NIH Office of Research Infrastructure Programs (P40 OD010440).

## Author contributions

PDM performed experiments and wrote the manuscript, JMD, DKH, BFH, JHX, and AMM performed experiments, DMR provided scientific guidance, and CFY supervised the project and edited the manuscript.

## Data availability

The datasets and strains generated during the current study are available from the corresponding author on reasonable request.

