## Supplementary Information for "A quiescent state following mild sensory arousal in *Caenorhabditis elegans* is potentiated by stress"

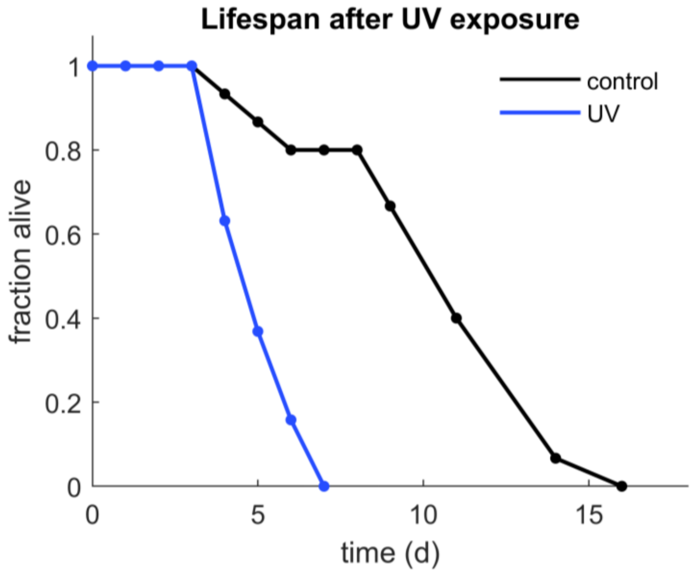


**Supplementary Figure S1. UV exposure reduces lifespan**. Lifespan of young adult WT animals after exposure to 1000 J/m$^2$ UV radiation (blue, n = 19) and unexposed same-chip controls (black, n = 15). Dots represent days when animals were scored. Mean lifespans: UV-treated 5.2 ± 0.3 days, control 10.9 ± 0.9 days, p = 0.00008, Wilcoxon rank-sum test.

### Supplementary Discussion 1. Similarity of the time course of UV SIS and EGF quiescence with and without mechanostimuli.

To study PRQ, we subjected animals in SIS or EGF-induced quiescence to mechanical or photic stimuli, typically once every 15 minutes. These stimuli elicit escape behavior and may be stressful themselves, so it is possible that they could alter overall timecourse or SIS and / or EGF-induced "baseline" quiescence. While we did not perform experiments designed to directly investigate this possibility (*i.e*. subjecting only a subset of animals on the same chip to stimuli in order to account for chip to chip variation), we did record UV SIS in WT animals as well as EGF-induced quiescence in WT and *ceh-17* or *aptf-1* mutant backgrounds in the absence of additional stimuli **(Supplementary Fig. S2)**. The time course of quiescence in these animals (left column, panels a, c, e, and g) was largely similar to the time course of baseline quiescence (quiescence excluding data from within 12 min of a stimulus) in equivalent experiments where mechanostimuli were applied (righthand column, panels b, d, f, and h). In WT animals, quiescence peaks in the second hour and then slowly decreases over the remainder of the recording period, but there are differences in the time course for *ceh-17* and *aptf-1* mutants. Therefore, while for some experiments we show the overall time course of SIS or EGF quiescence using the baseline quiescence from experiments with stimuli (**Supplementary Figs. S5, S6, S7, and S10**), we cannot rule out an effect of the stimuli on the overall quiescence time course in these cases. In addition to these panels, we have shown the baseline quiescence from all other PRQ experiments where the baseline quiescence is not otherwise shown **(Supplementary Fig. S3)**.

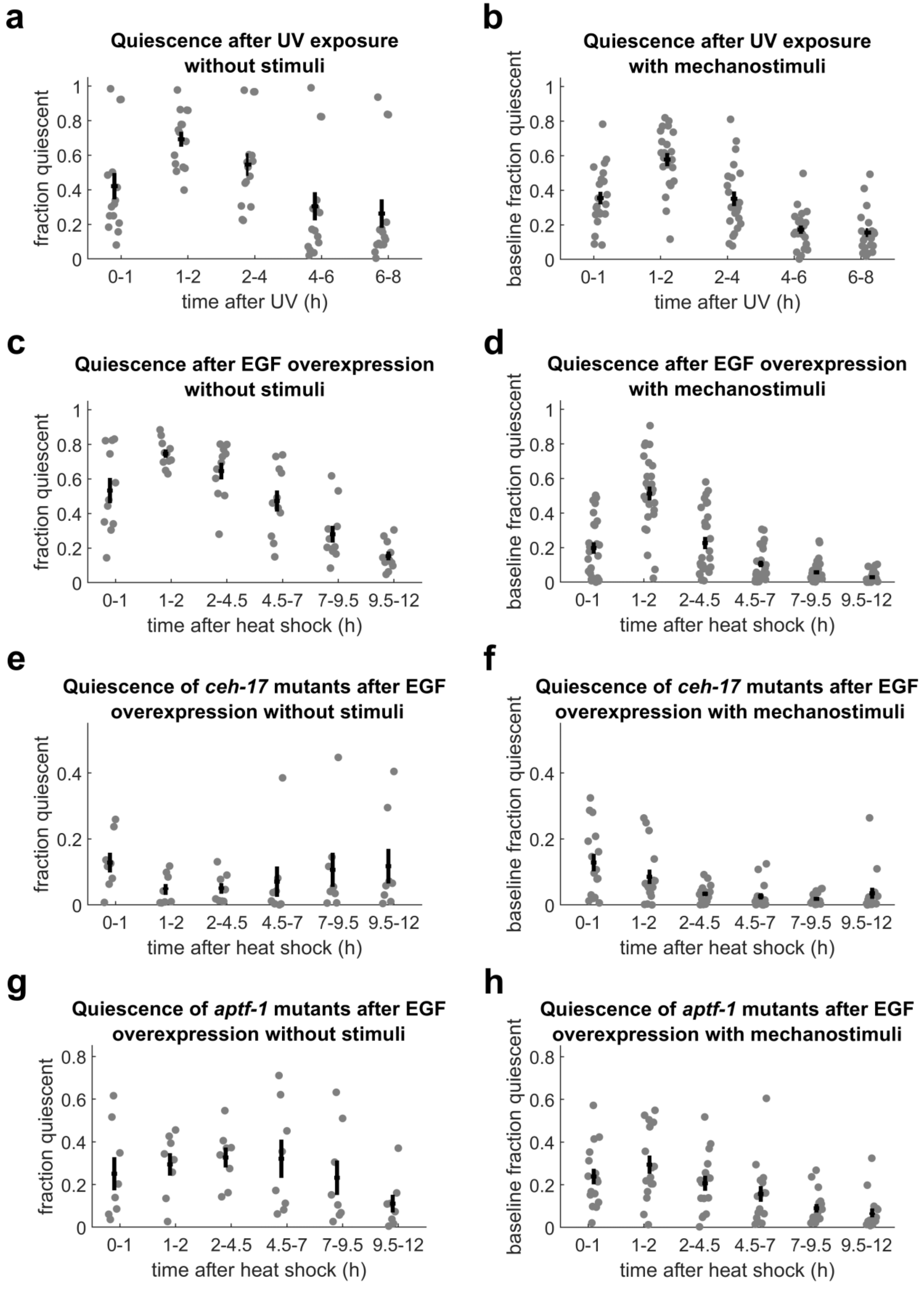
a
Quiescence after UV exposure without stimuli
fraction quiescent
time after UV (h)
0-1
1-2
2-4
4-6
6-8
b
Quiescence after UV exposure with mechanostimuli
baseline fraction quiescent
time after UV (h)
c
Quiescence after EGF overexpression without stimuli
fraction quiescent
time after heat shock (h)
0-1
1-2
2-4.5
4.5-7
7-9.5
9.5-12
d
Quiescence after EGF overexpression with mechanostimuli
baseline fraction quiescent
time after heat shock (h)
e
Quiescence of ceh-17 mutants after EGF overexpression without stimuli
fraction quiescent
time after heat shock (h)
f
Quiescence of ceh-17 mutants after EGF overexpression with mechanostimuli
baseline fraction quiescent
time after heat shock (h)
g
Quiescence of aptf-1 mutants after EGF overexpression without stimuli
fraction quiescent
time after heat shock (h)
h
Quiescence of aptf-1 mutants after EGF overexpression with mechanostimuli
baseline fraction quiescent
time after heat shock (h)

**Supplementary Figure S2. Time course of UV SIS and EGF quiescence with and without mechanostimuli.**

**(a)** Fraction of time quiescent of WT animals after 1000 $J/m^2$ UV and in the absence of stimuli. n = 15 animals, four replicates. These data also appear in Fig. 1.

**(b)** Baseline fraction of time quiescent of WT animals after 1000 $J/m^2$ UV and in the presence of mechanostimuli. n = 22 animals, two replicates. These data also appear in Fig. 1.

**(c)** Fraction of time quiescent of WT background animals after a mild, 10 min, 32 °C, heat shock in the absence of stimuli. n = 11 animals, two replicates. These data also appear in Fig. 2.

**(d)** Baseline fraction of time quiescent of WT background animals after a mild, 10 min, 32 °C, heat shock in the presence of mechanostimuli. n = 28 animals, four replicates. These data also appear in Fig. 2.

**(e)** Fraction of time quiescent of *ceh-17* mutant background animals after a mild, 10 min, 32 °C, heat shock in the absence of stimuli. n = 8 animals, one replicate. These data also appear in Fig. 4.

**(f)** Baseline fraction of time quiescent of *ceh-17* mutant background animals after a mild, 10 min, 32 °C, heat shock in the presence of mechanostimuli. n = 16 animals, two replicates. These data also appear in Fig. 4.

**(g)** Fraction of time quiescent of *aptf-1* mutant background animals after a mild, 10 min, 32 °C, heat shock in the absence of stimuli. n = 8 animals, one replicate. These data also appear in Fig. 4.

**(h)** Baseline fraction of time quiescent of *aptf-1* mutant background animals after a mild, 10 min, 32 °C, heat shock in the presence of mechanostimuli. n = 16 animals, two replicates. These data also appear in Fig. 4.

Dots in all panels represent individual animal averages in the specified time bins. For all experiments with stimuli, data recorded within 12 min of a stimulus were excluded from analysis.

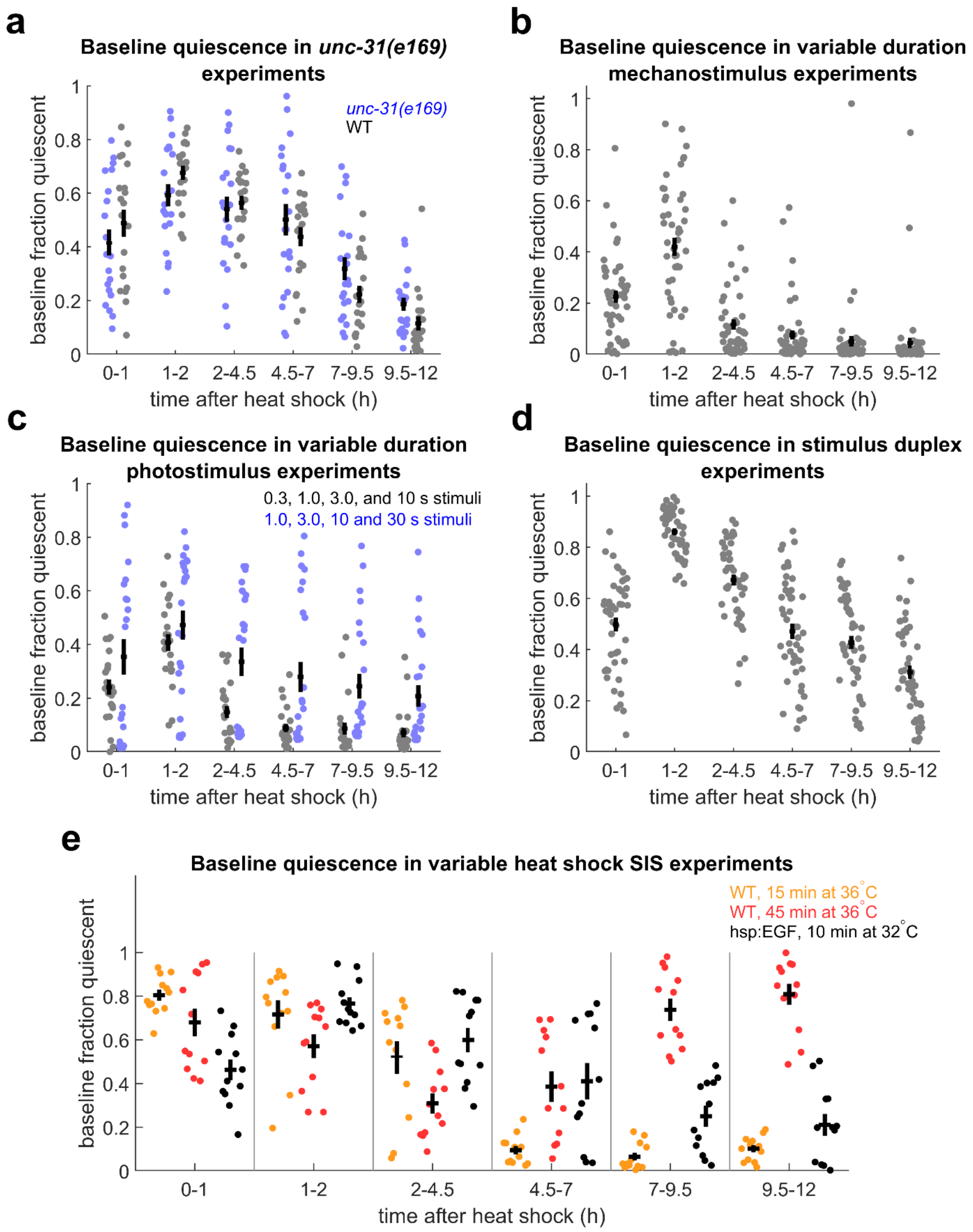
a
Baseline quiescence in unc-31(e169) experiments
unc-31(e169)
WT
baseline fraction quiescent
time after heat shock (h)
0-1
1-2
2-4.5
4.5-7
7-9.5
9.5-12
b
Baseline quiescence in variable duration mechanostimulus experiments
c
Baseline quiescence in variable duration photostimulus experiments
0.3, 1.0, 3.0, and 10 s stimuli
1.0, 3.0, 10 and 30 s stimuli
d
Baseline quiescence in stimulus duplex experiments
e
Baseline quiescence in variable heat shock SIS experiments
WT, 15 min at 36°C
WT, 45 min at 36°C
hsp:EGF, 10 min at 32°C

**Supplementary Figure S3. Time course of baseline quiescence after EGF overexpression in PRQ experiments.**

**(a)** Baseline fraction of time quiescent of *unc-31(e169)* mutant and WT background animals following EGF overexpression. n = 21 *unc-31(e169)* and 20 WT background animals from two replicates. Data from the same experiments appear in Supplementary Fig. S9.
**(b)** Baseline fraction of time quiescent for WT animals following EGF overexpression in the presence of mechanostimuli of varying duration. n = 48 animals, four replicates. Data from these experiments appear in Fig. 5b and supplementary Fig. S12a.
**(c)** Baseline fraction of time quiescent for WT animals following EGF overexpression in the presence of photostimuli of varying duration. n = 23 animals, two replicates. Data from these experiments appear in Fig. 5c and Supplementary Fig. S12b.
**(d)** Baseline fraction of time quiescent for WT animals following EGF overexpression in the single / duplex stimulus experiments shown in Fig. 5. n = 47 animals, two replicates.
**(e)** Baseline fraction of time quiescent for WT animals following heat shock and EGF overexpression controls from the variable heat shock experiment shown in Supplementary Fig. S4. n = 12 animals, two replicates each condition.
In all panels, baseline fraction (of time) quiescent was calculated excluding data recorded within 12 min of a stimulus.

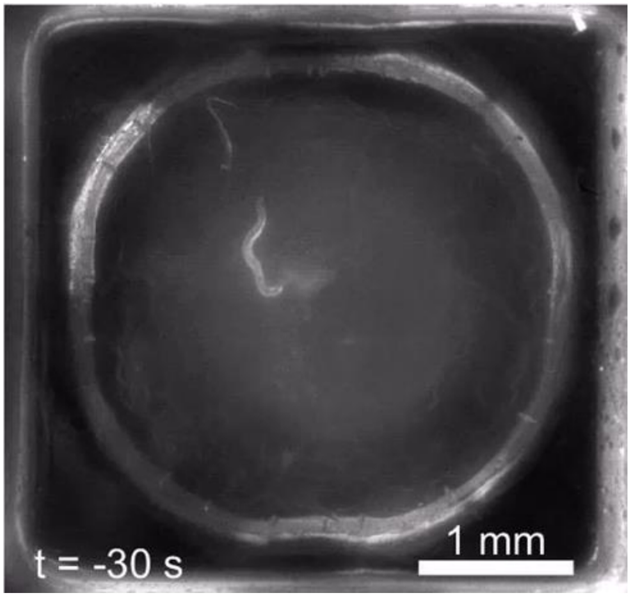


**Supplementary Video 1. Post-response quiescence in a WT animal approximately 4 h after exposure to 1000 J/m$^2$ UV-C radiation.** The stimulus is a 1 kHz, 1 s substrate vibration. Time is displayed relative to the timing of the mechanical stimulus. Original video is 2 min duration at 1 fps.

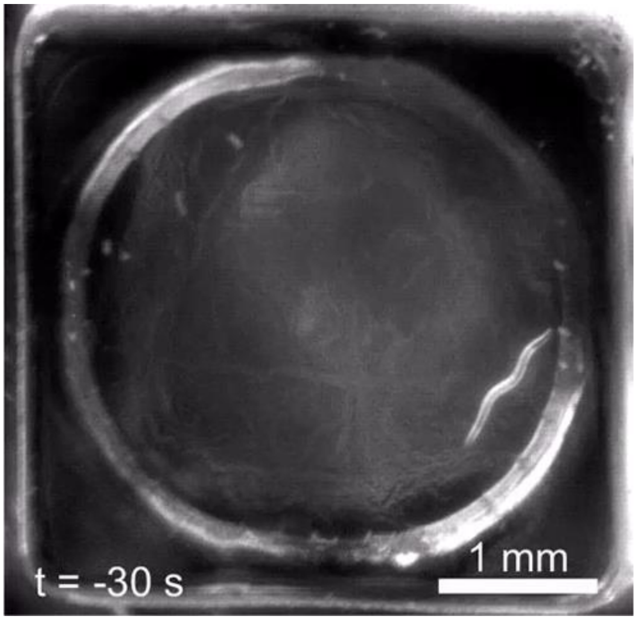


**Supplementary Video 2. Normal mechanosensory response of a non UV-treated animal.** There is no quiescence following the locomotor response and turn. The stimulus is a 1 kHz, 1 s substrate vibration. Time is given relative to the timing of the mechanical stimulus. Original video is 2 min duration at 1 fps.

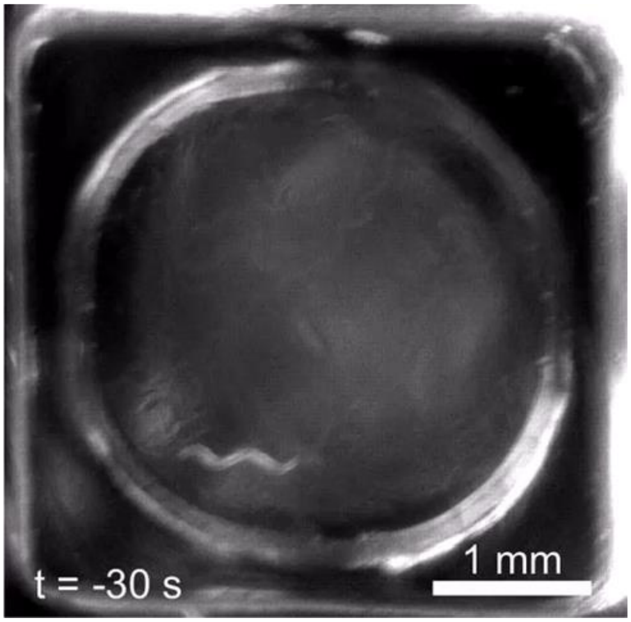


**Supplementary Video 3. Post-response quiescence following mechanosensory stimulus in a non UV control animal.** The stimulus is a 1 kHz, 1 s substrate vibration. Time is given relative to the timing of the mechanical stimulus. Original video is 2 min duration at 1 fps.

**Supplementary Discussion 2: PRQ does not arise from synchronization of active or quiescent bouts.**

SIS and EGF quiescence consist of alternating active and quiescent bouts. Therefore, it is possible that PRQ, a transient increase in quiescent behavior observed after a stimulus, is the result of temporary synchronization of quiescent bouts following the stimulus. To test this hypothesis, we investigated three different methods of aligning baseline (pre-stimulus) quiescence data (binary active / quiescent scores of each animal at every frame) of UV SIS **(Supplementary Fig. S4a)** and EGF overexpressing **(Supplementary Fig. S4b)** animals and compared the resulting average quiescence traces to actual PRQ in the same animals.

Because the mechanosensory response begins with a bout of activity, we first aligned the baseline quiescence data by the beginning of spontaneous active bouts. The resulting average quiescence trace (labeled “active bouts aligned”) shows a trough in quiescence followed by a return to baseline, rather than a PRQ-like peak in quiescence.

Next, we considered the possibility that the stimulus causes a stereotyped bout of activity associated with the mechanosensory response which serves to align the subsequent quiescent bouts. To simulate this scenario, we first aligned baseline quiescence data by the beginning of quiescent bouts, but then modified the baseline quiescence data by inserting a period of activity equal to the duration of each animal's initial active response to the next stimulus **(Supplementary Fig. S4c)**. For example, if an animal became continuously active for 13 s following the next stimulus, we would insert 13 s of activity into its baseline quiescence data, pushing the baseline data after the insertion point forward in time. The resulting average quiescence traces (labeled “quiescent bouts aligned, offset by response”) consist of a decrease in quiescence associated with the inserted response activity periods followed by a very small peak in quiescence of a magnitude much lower than actual PRQ.

Lastly, we simulated the perfect synchronization of quiescent bouts by aligning the baseline quiescence data by the beginning of quiescent bouts. The resulting average quiescence trace (labeled “quiescent bouts aligned”) shows a sharp peak in quiescence which is of greater amplitude and decays much more quickly than PRQ. The area under this peak appears much smaller than that under the PRQ peak, implying that PRQ cannot be recapitulated by synchronizing normal SIS behavior.

Taken together, these results indicate that PRQ is not caused by the synchronization of otherwise normal SIS or EGF-induced quiescent bouts.

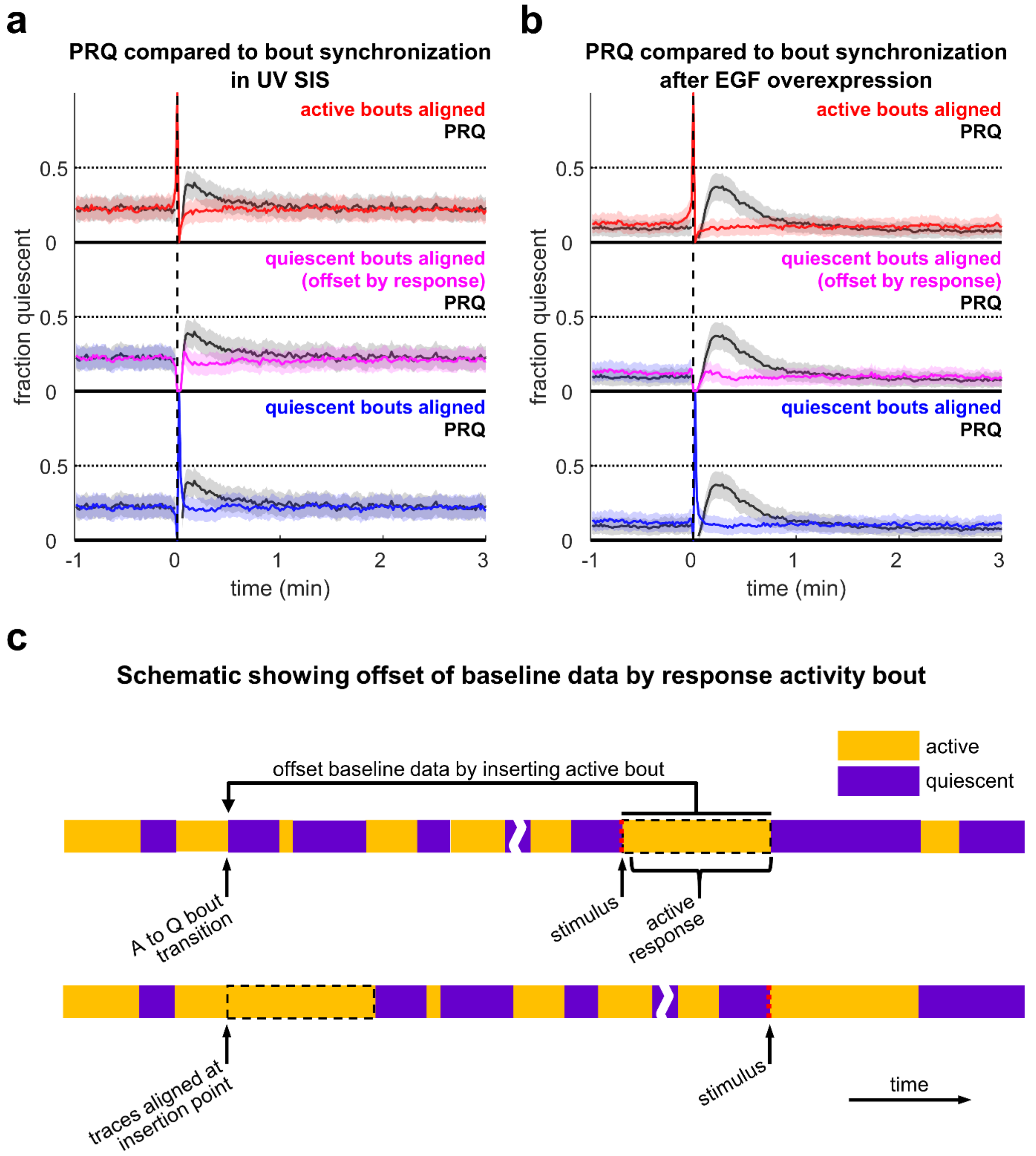


**Supplementary Figure S4. Comparison of computational alignment of SIS quiescent or active bouts with quiescence dynamics during PRQ.**
**(a) (top)** Fraction quiescent of WT animals during UV SIS following mechanosensory stimulus (black) or *in silico* alignment of active bouts (red). **(middle)** Fraction quiescent of the same animals following a mechanosensory stimulus (black) or *in silico* alignment of quiescent bouts offset by the duration of the first active period following the next stimulus for that animal (magenta). **(bottom)** Fraction quiescent of the same animals following a mechanosensory stimulus (dashed line) or *in silico* alignment of quiescent traces (blue). n = 22 animals, two replicates. Data from the same experiment are shown in Fig. 1a-c.

**(b) (top)** Fraction quiescent of WT background animals after EGF overexpression following a mechanosensory stimulus (black) or *in silico* alignment of active traces (red). **(middle)** Fraction quiescent of the same animals following a mechanosensory stimulus (black) or *in silico* alignment of quiescent traces offset by the duration of the first active period following the next stimulus for that animal (magenta). **(bottom)** Fraction quiescent of the same animals following a mechanosensory stimulus (black) or *in silico* alignment of quiescent traces (blue). n = 28 animals, four replicates. Data from the same experiment are shown in Fig. 2a-c.
**(c)** Schematic showing the method used to offset baseline quiescence data to create the middle (magenta) traces in panels (a) and (b). A fictitious activity (A, yellow) and quiescence (Q, purple) trace from a single animal is shown. An active bout equal in duration to the first active response bout after the stimulus is inserted into baseline data at an active to quiescent bout transition. Binary traces created this way and aligned by the insertion point and averaged to create the traces in panels (a) and (b), middle. The break in the timeline represents the passage of time (at least three minutes) between the insertion area (left) and the real stimulus (right).

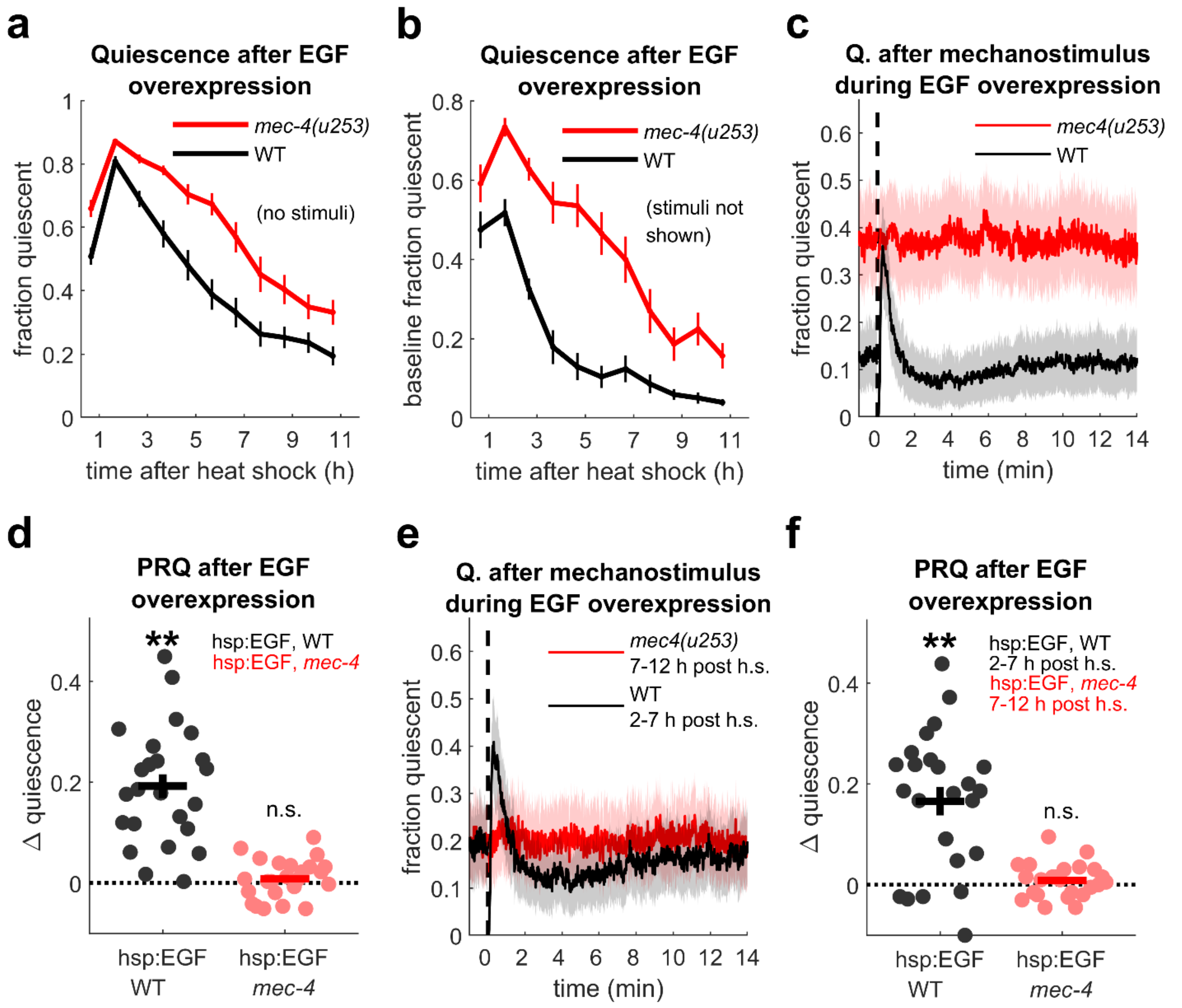


**Supplementary Figure S5. Mechanostimulus does not affect quiescence in a *mec-4* loss of function mutant.**

**(a)** Fraction of time quiescent of *mec-4(u253)* and WT background animals following EGF overexpression in the absence of stimuli. n = 24 animals per condition, two replicates. Traces show fraction of time quiescent ± SEM over 60 min.

**(b)** Baseline fraction of time quiescent of *mec-4(u253)* and WT background animals following EGF overexpression, n = 24 animals per condition, two replicates. Traces show fraction of time quiescent ± SEM over 60 min, excluding the 12 min following mechanostimuli, which are not shown.

**(c)** Quiescence of *mec-4* mutant (red) and WT (black) background animals before and after 1kHz, 1 s vibrations (dashed line), 2-12 h after EGF overexpression (same animals as in panel (a)). Shading represents ± SEM.

**(d)** Quantification of PRQ for data in panels (a) and (b); dots represent single animal averages for stimuli 2-12 h after EGF overexpression. ** denotes significance at α = 0.01 (one sample, two-tailed t-test with Bonferroni correction for two comparisons). Error bars show ± SEM.

**(e)** Data from the same experiment as panel (b) except only data from 7-12 h post shock for *mec-4* animals, and 2-7 h post heat shock for WT-background animals, is shown. Baseline quiescence was similar between the two strains during these time periods.
**(f)** Quantification of PRQ for data restricted as described in panel (d). ** denotes significance at α = 0.01 (one sample, two-tailed t-test with Bonferroni correction for two comparisons).
∆ quiescence was calculated as in Fig. 1a.

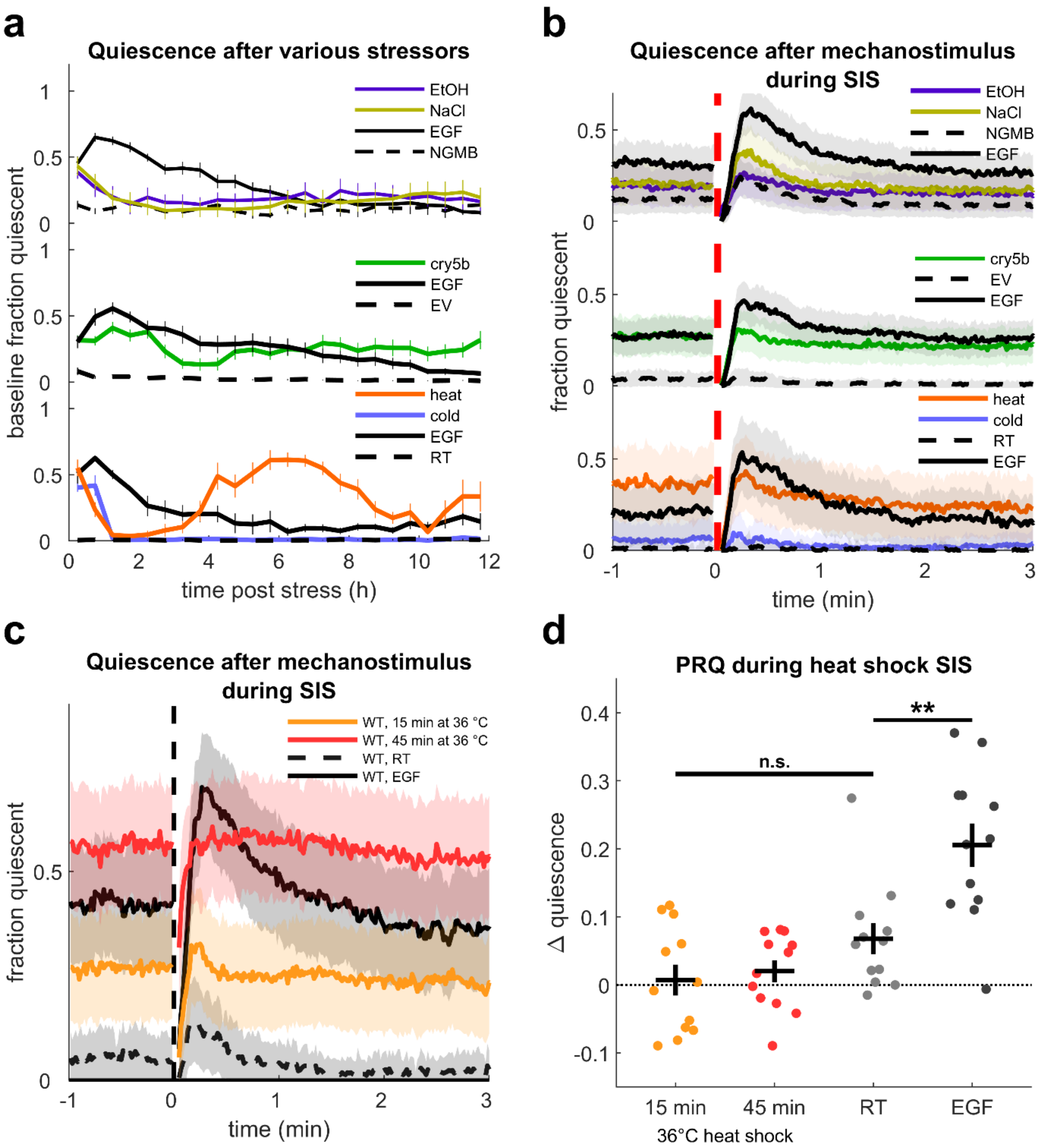


**Supplementary Figure S6. Baseline quiescence and quiescence after mechanosensory stimulus following exposure to various stressors.**
**(a**) **(top)** Baseline fraction of time quiescent of WT animals following 15 min immersion in 5% EtOH in NGMB, NGMB with an added 500 mM NaCl, or standard NGMB, and EGF overexpression (three replicates, 12 to 18 animals per condition); **(middle)** WT animals exposed

to Cry5b-expressing bacteria for 15 min, WT animals exposed to empty vector (EV) control bacteria for 15 min, and EGF overexpression (three replicates, 17 to 22 animals per condition); and **(bottom)** WT animals following exposure to 36°C for 30 min, 4 °C for 24 h, room temperature controls compared to EGF overexpression (two replicates, 12 animals per condition). Data are the same as in Fig. 3. Stimuli are not shown, and data recorded within 12 min of a stimulus were excluded from analysis. Error bars show ± SEM.

**(b)** Movement quiescence before and after 1 s, 1 kHz substrate vibrations (dashed line) after stressors that cause SIS. The inter-stimulus interval was 15 min. Data are from the same experiments as panel (a) and Fig. 3. Shading shows ± SEM.

**(c)** Movement quiescence before and after 1 s, 1 kHz substrate vibrations (vertical dashed line) following 15 min (orange) and 45 min (red) heat shock as well as room temperature (dashed black) and EGF overexpression (solid black). The inter-stimulus interval was 15 min. Traces represent 12 animals per condition from two replicates. Shading shows ± SEM.

**(d)** Quantification of PRQ for data in panel (c). Dots represent single animal averages for stimuli 0-12 h after heat shock or EGF overexpression or in controls held at room temperature. $\Delta$ quiescence was calculated as in Fig. 1a. ** denotes significance at $\alpha = 0.01$ (two sample, two-tailed t-test with Bonferroni correction for three comparisons). Error bars show ± SEM.

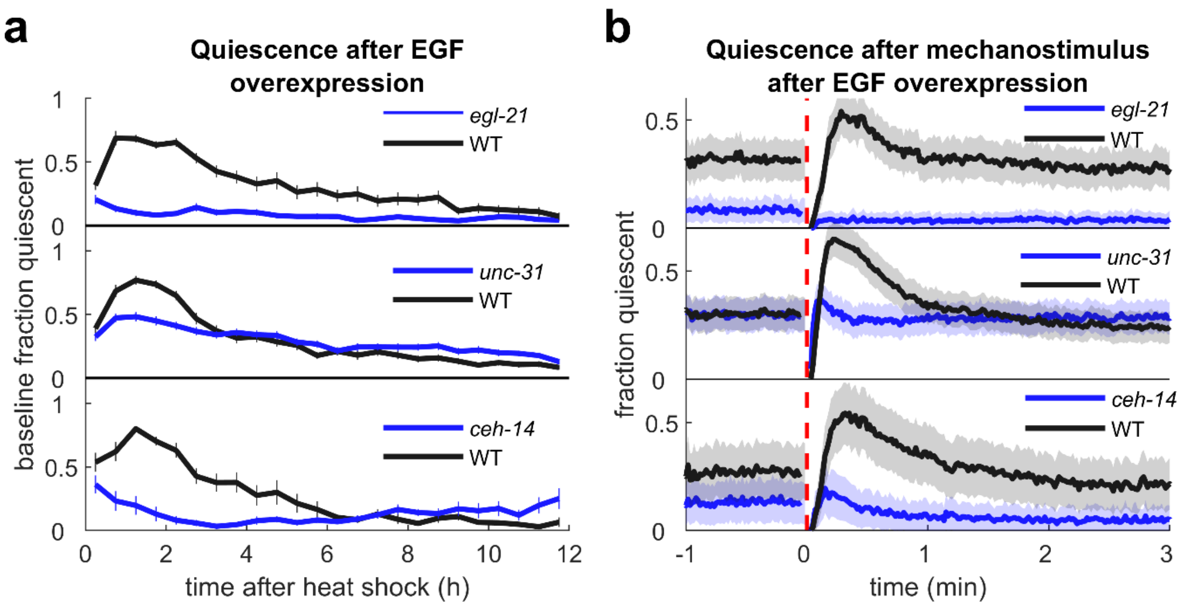


**Supplementary Figure S7. Overall and post stimulus quiescence of *egl-21*, *unc-31*, and *ceh-14* mutants after EGF overexpression.**

**(a)** 30 min average baseline fraction of time quiescent of WT (black) and *egl-21* **(top)**, *unc-31* **(middle)**, or *ceh-14* **(bottom)** mutants (blue) for 12 h following EGF overexpression. Data from the 12 min after mechanostimuli are excluded from the average. Stimuli not shown. n = 24 animals, two replicates per strain for *egl-21* experiments; n = 36 mutant and 35 WT, three replicates for *unc-31* experiments; n = 12 animals, one replicate per strain for *ceh-14* experiments. Data from the same experiment appear in Fig. 4c. Error bars show ± SEM.

**(b) F**raction quiescent before and after mechanostimulus (dashed red line) in WT (black) and mutant (blue) background animals 2-12 h following EGF overexpression. Data are from the same experiment as panel (a) and Fig. 4c. Shading shows ± SEM.

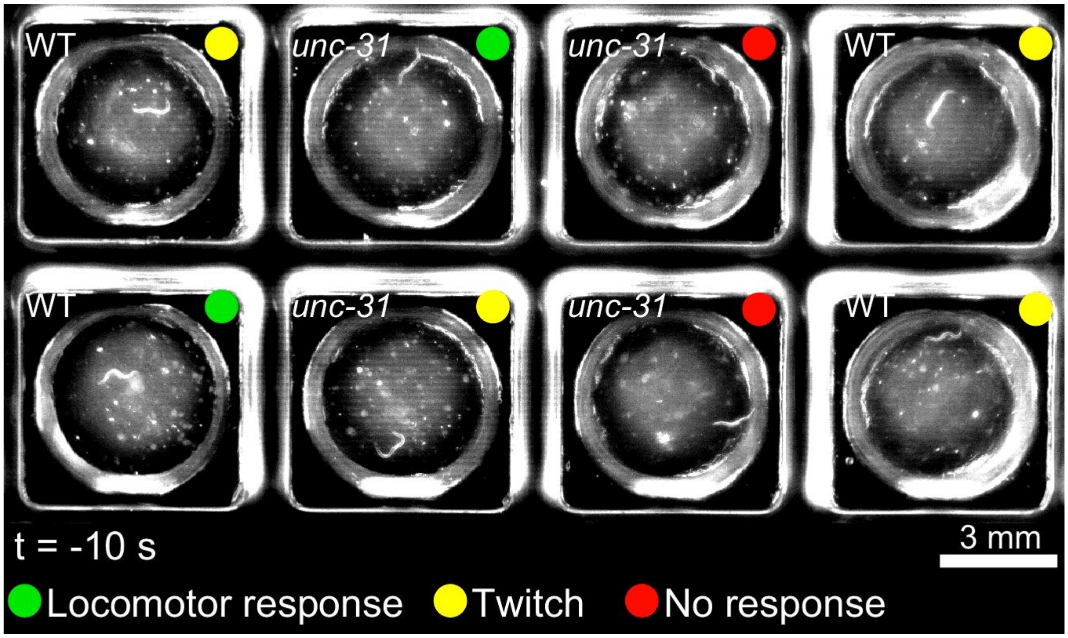


**Supplementary Video 4. Mechanosensory responses of WT and *unc-31* mutants.**
Visual inspection of the mechanosensory response to substrate vibrations reveals that while some animals (green dots) move after the stimulus, others (yellow dots) twitch or move only during the stimulus and then remain quiescent, and some *unc-31* animals (red dots) do not appear to respond. Slight movement of the wormotel during mechanostimulus complicated quantification of twitch responses using difference images from consecutive frames, so twitch responses were detected by subtracting non-consecutive frames that straddled the stimulus **(Supplementary Figs. S8, S9)**.

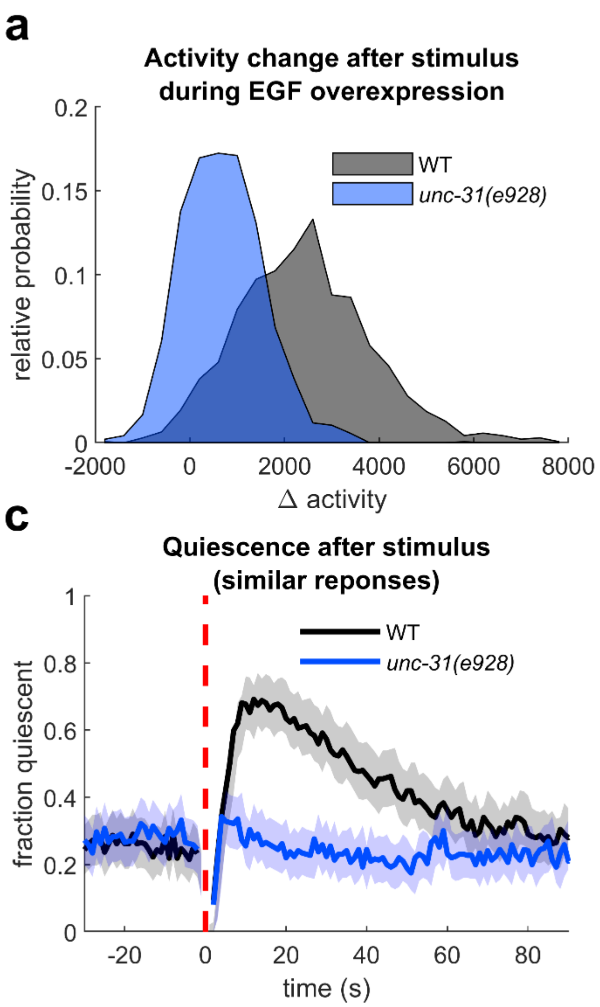


**Supplementary Figure S8.** ***unc-31(e928)*** **mutants are defective in PRQ even after controlling for the magnitude of mechanosensory response.**

*unc-31* mutant animals are less responsive to substrate vibrations, and sometimes do not respond to these stimuli. Therefore, they may lack PRQ simply because they lack a response. To control for this, we measured the responses of WT and *unc-31(e928)* animals and examined the change in quiescence following stimuli in animals with a similar response magnitude. We used non-consecutive frame subtraction to detect "twitch" type responses wherein all movement takes place during the stimulus (see Supplementary Vid. 4). We found that *unc-31* animals appear to be defective in PRQ compared to WT background animals even when controlling for response magnitude.

**(a)** Histogram of the mechanosensory responses of WT and *unc-31(e928)* background animals 2-12 h after EGF overexpression. Stimuli were 1 s, 1 kHz substrate vibrations, responses are activity in the 5 s straddling the stimulus minus the mean activity in the 20 s preceding the stimulus. (n = 36 *unc-31(e928)* and 35 WT animals from three replicates and a total of 2840 responses). Data from these experiments also appear in Fig. 4c.

**(b)** Average fraction quiescent before and after all stimuli (dashed red line) of *unc-31(e928)* (blue) and WT (black) background animals carrying the hsp:EGF array 2-12 h after mild heat shock. Traces are the averages of 1440 *unc-31(e928)* and 1400 WT responses from the same animals as panel (a). Shading in panels b-d show ± SEM.

**(c)** Average fraction quiescent before and after stimuli (dashed red line) to which responses fell between 1200 and 2000 activity values, 2-12 h after mild heat shock. Traces are the averages of 286 *unc-31(e928)* and 279 WT responses from the same animals as panel (a).
**(d)** Average fraction quiescent before and after stimuli (dashed red line) to which responses were $\leqslant 0$ activity values, 2-12 h after mild heat shock. Traces are the averages of 318 *unc-31(e928)* and 43 WT responses from the same animals as panel (a).

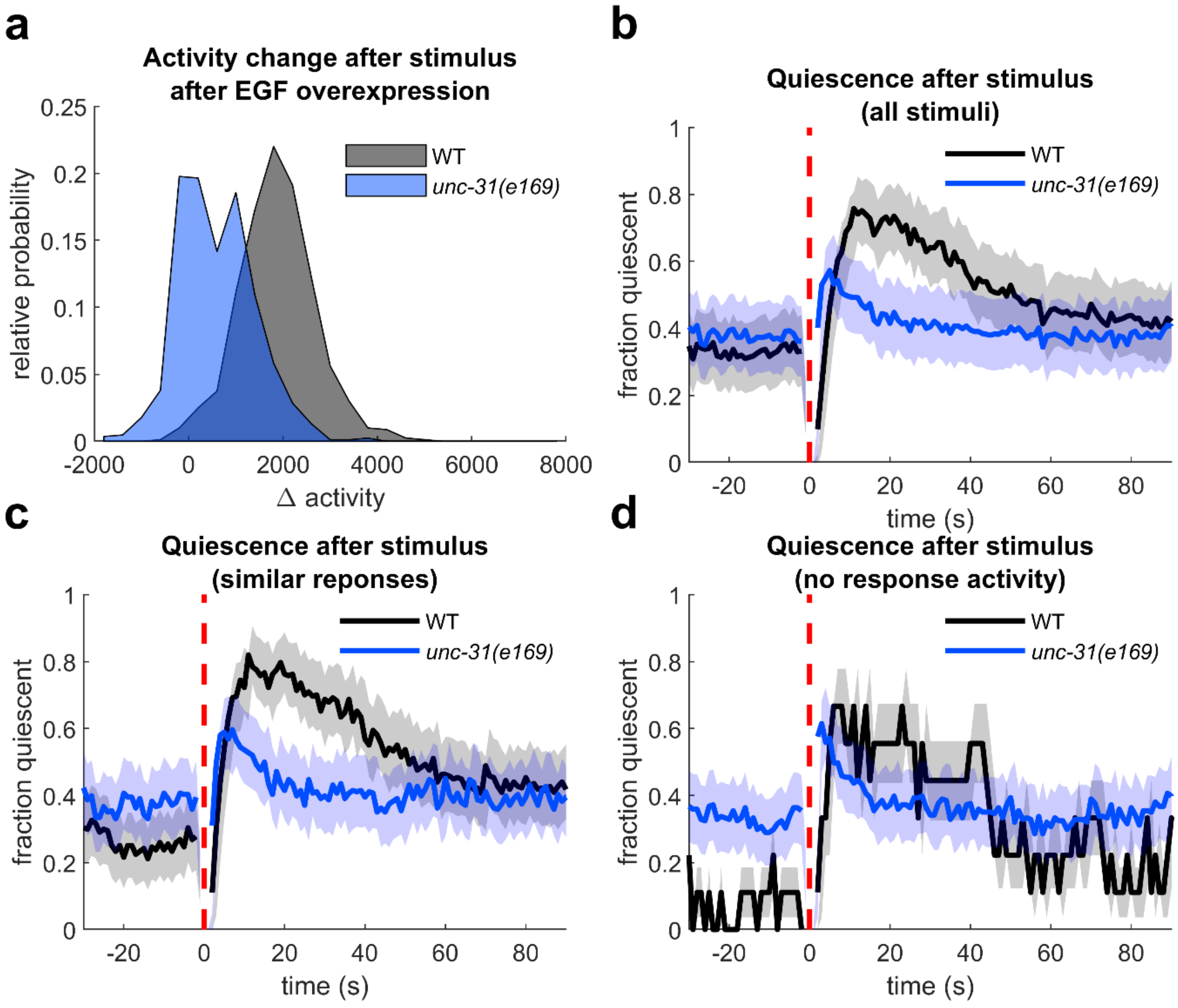


**Supplementary Figure S9. *unc-31(e169)* mutants are partially defective in PRQ even after controlling for the magnitude of mechanosensory response.**

*unc-31* mutant animals are less responsive to substrate vibrations, and sometimes do not respond to these stimuli. In this figure we compare pre- and post-stimulus quiescence in animals with similar amounts of response activity. The experiments and analysis in this figure are equivalent to Supplementary Fig. S8, but *unc-31(e169)* is substituted for *unc-31(e928*).

**(a)** Histogram of the responses of WT and *unc-31(e169)* background animals 2-12 h after EGF overexpression. Stimuli were 1 s, 1 kHz substrate vibrations, responses are activity in the 5 s straddling the stimulus minus the mean activity in the 20 s preceding the stimulus. (n = 21 *unc-31(e169)* and 20 WT background animals from two replicates and a total of 1640 responses).

**(b)** Average fraction quiescent before and after *all* stimuli (dashed red line) of *unc-31(e169)* (blue) and WT (black) background animals carrying the hsp:EGF array 2-12 h after mild heat shock. Traces are the averages of 840 *unc-31* and 800 WT responses from the same animals as panel *a*. Shading in panels b-d show ± SEM.

**(c)** Average fraction quiescent before and after stimuli (dashed red line) after which responses fell between 800 and 1600 activity values, 2-12 h after mild heat shock. Traces are the averages of 248 *unc-31* and 227 WT responses from the same animals as panel (a).

**(d)** Average fraction quiescent before and after stimuli (dashed red line) to which responses were $\leqslant 0$ activity values, 2-12 h after mild heat shock. Traces are the averages of 231 *unc-31* and 9 WT responses from the same animals as panel (a).

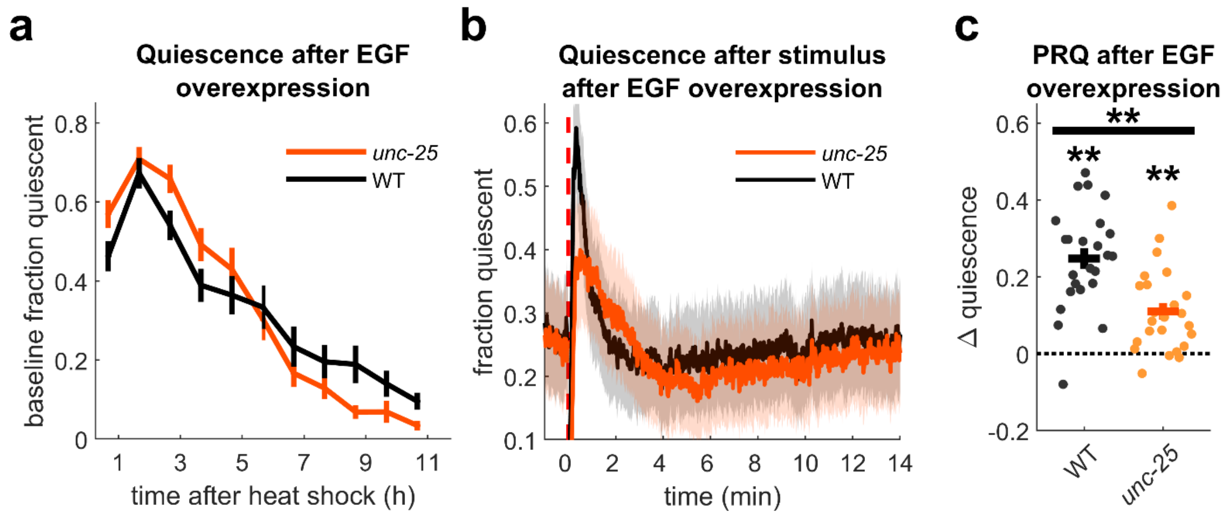


**Supplementary Figure S10. Loss of function of *unc-25* mutants have lower peak PRQ.**
**(a)** Baseline fraction of time quiescent of WT and *unc-25(e156)* animals following EGF overexpression. Traces show average quiescence over 60 min, excluding the 12 min following each mechanostimulus n = 24 animals per strain, two replicates. Error bars here and in panel (c) indicate ± SEM.
**(b)** Quiescence of WT (black) and *unc-25* (orange) animals before and after a 1 kHz, 1 s mechanostimulus, same animals as in panel (a). Shading indicates ± SEM.
**(c)** Quantification of PRQ from panel (b). Change in quiescence (Δ quiescence) is calculated as in Fig. 1a. Dots represent single animal averages across all stimuli. ** denotes $p < 0.01$, two-tailed t-test (testing for significant PRQ in each strain), or two sample t-test (comparing PRQ of WT and *unc-25* background animals).

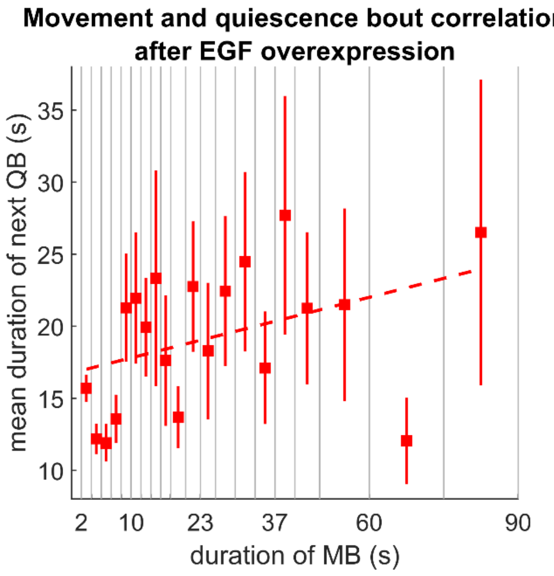


**Supplementary Figure S11. Correlation of the duration of quiescent bouts (QBs) with the preceding movement bout (MB) in EGF quiescence.**
Squares represent mean QB duration following MBs in each duration bin with bounds at 2, 4, 6, 8, 10, 12, 14, 16, 18, 20, 23, 26, 29, 33, 37, 41, 45, 50, 60, 75, and 90 s, bars represent SEM. Bout duration was positively but not significantly correlated. (slope = 0.088, $p = 0.089$, $R^2 = 0.15$). n = 12 animals, two replicates.

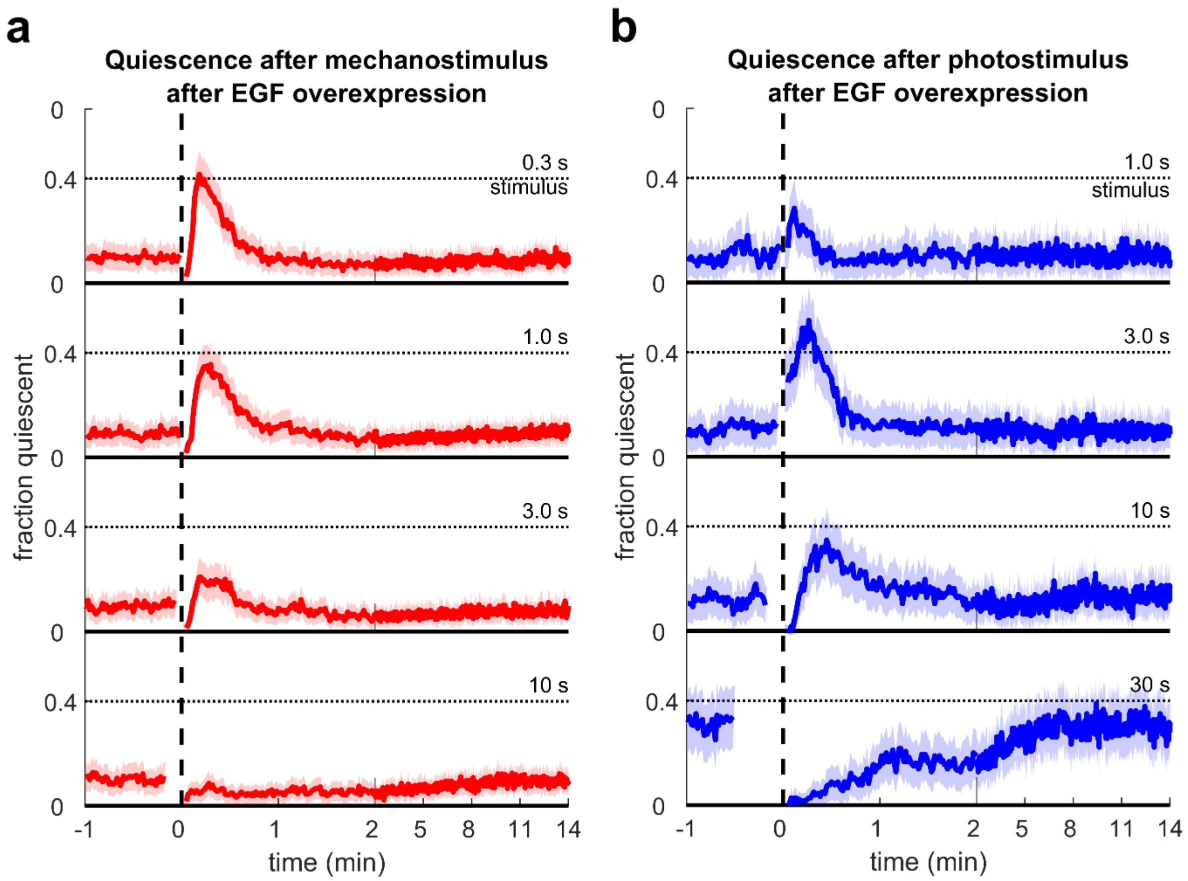


**Supplementary Figure S12. Fraction quiescent following mechanosensory and blue light stimuli of various durations.**
**(a)** Fraction quiescent of WT background animals 2-12 h after EGF overexpression following 0.3, 1.0, 3.0, and 10 s duration mechanosensory stimuli. n = 48 animals, four replicates. Shading here and in panel (b) indicates ±SEM.
**(b)** Fraction quiescent of WT background animals 2-12 h after EGF overexpression following 1.0, 3.0, 10, and 30 s duration blue light stimuli. n = 23, two replicates for 1, 3, and 10 s, n = 24, three replicates, for 30 s stimuli.

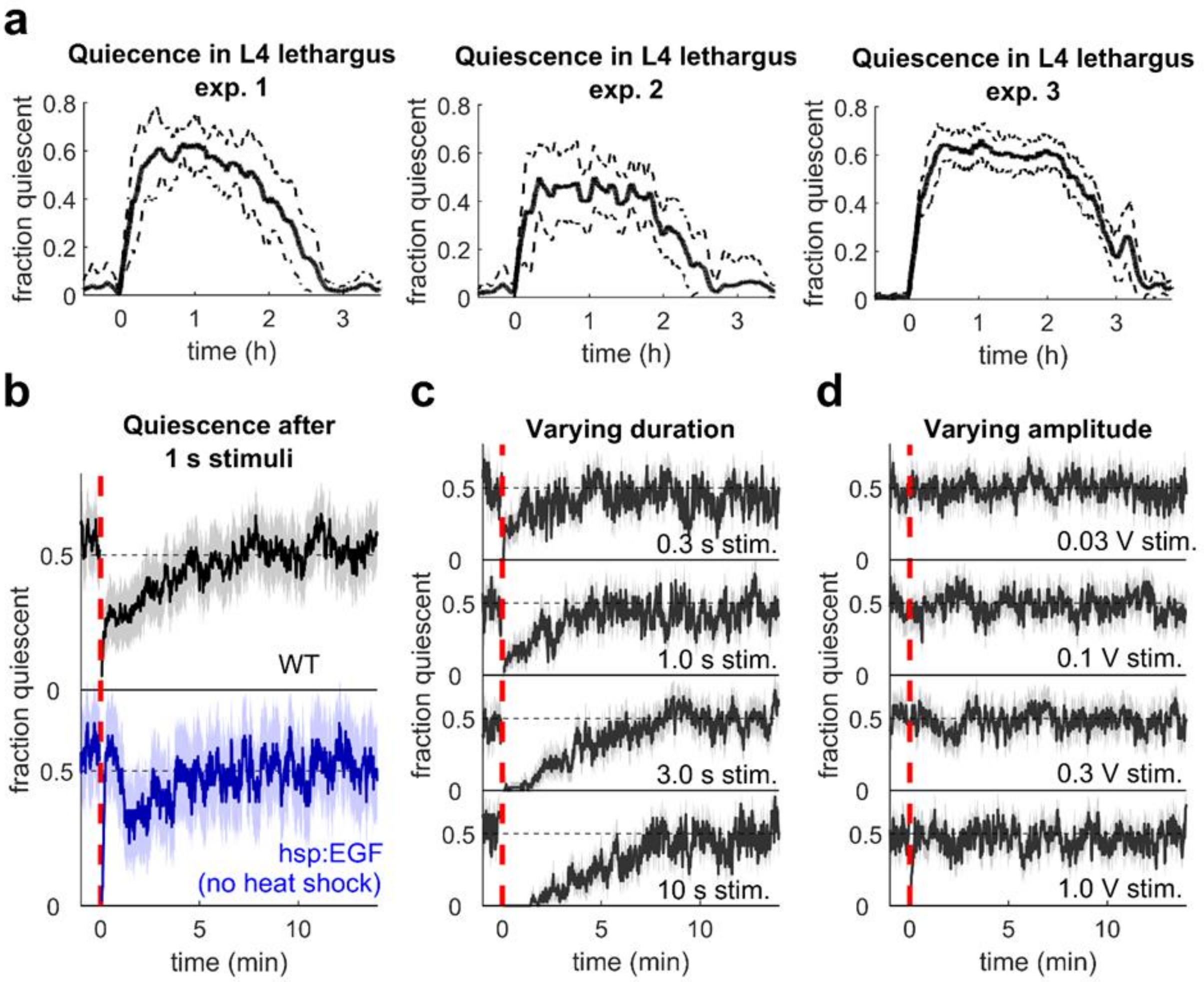


**Supplementary Figure S13. Post response quiescence does not occur after mechanostimulus during L4 lethargus.**

We observed PRQ after 0.3 and 1.0 s substrate vibrations following EGF overexpression (Fig. 5b). A similar increase in quiescence has been reported following short duration (1 s) but not long duration (15 s) 1 kHz substrate vibrations separated by a 15 min ISI during lethargus (Nagy *et al*. 2014). We performed three experiments to attempt to replicate this finding. We staged L4 animals by vulva morphology and active feeding and restricted analysis to lethargus as defined previously (see Raizen *et al.* 2008 and Iwanir *et al.* 2013). Briefly, after smoothing with a 10 min filter, the beginning of lethargus was defined as the point after which quiescence levels stayed above 5% for at least 20 min, and the end of lethargus was defined as the point after which quiescence levels stayed below 5% for at least 20 min. We performed three experiments (1) We stimulated WT and hsp:EGF (no heat shock) lethargus animals every 15 min with 1 s 1 kHz vibrations, (2) we stimulated WT lethargus animals every 15 min with 0.3, 1.0, 3.0, and 10 s, 1 kHz substrate vibrations, and (3) we stimulated WT lethargus animals with 0.3 s, 1 kHz substrate vibrations, but changed the amplitude of the audio signal sent to the amplifier from 1 V (used in all other experiments in this manuscript) to 0.3, 0.1, and 0.03 V, zero to peak. Each experiment had one replicate. We plotted the quiescence of lethargus animals after these stimuli; in no case did we observe an increase in quiescence above baseline.

**(a)** Fraction of animals quiescent during lethargus for experiments 1, 2, and 3 plotted against time after lethargus detected for each animal. Dashed lines indicate ± standard deviation.

Mechanostimuli are present but not shown, and data from the frames immediately prior to a stimulus to the frames immediately (three frames per stimulus) after are excluded from this plot

**(b)** Average quiescence of WT (black, n = 16) and hsp:EGF (blue, n = 8) lethargus animals following 1 s, 1 kHz substrate vibrations (red dashed line). Animals were not heat shocked; the brief recovery of baseline quiescence of hsp:EGF animals may be due to leaky expression of EGF. n = 10 (WT), 6 (hsp:EGF). Shading in panels b-d indicates ± SEM.

**(c)** Average quiescence of WT lethargus animals following 1 kHz substrate vibrations of various durations (red dashed line). n = 16.

**(d)** Average quiescence of WT lethargus animals following 1 kHz, 0.3 s substrate vibrations of various amplitudes (red dashed line). n = 17.

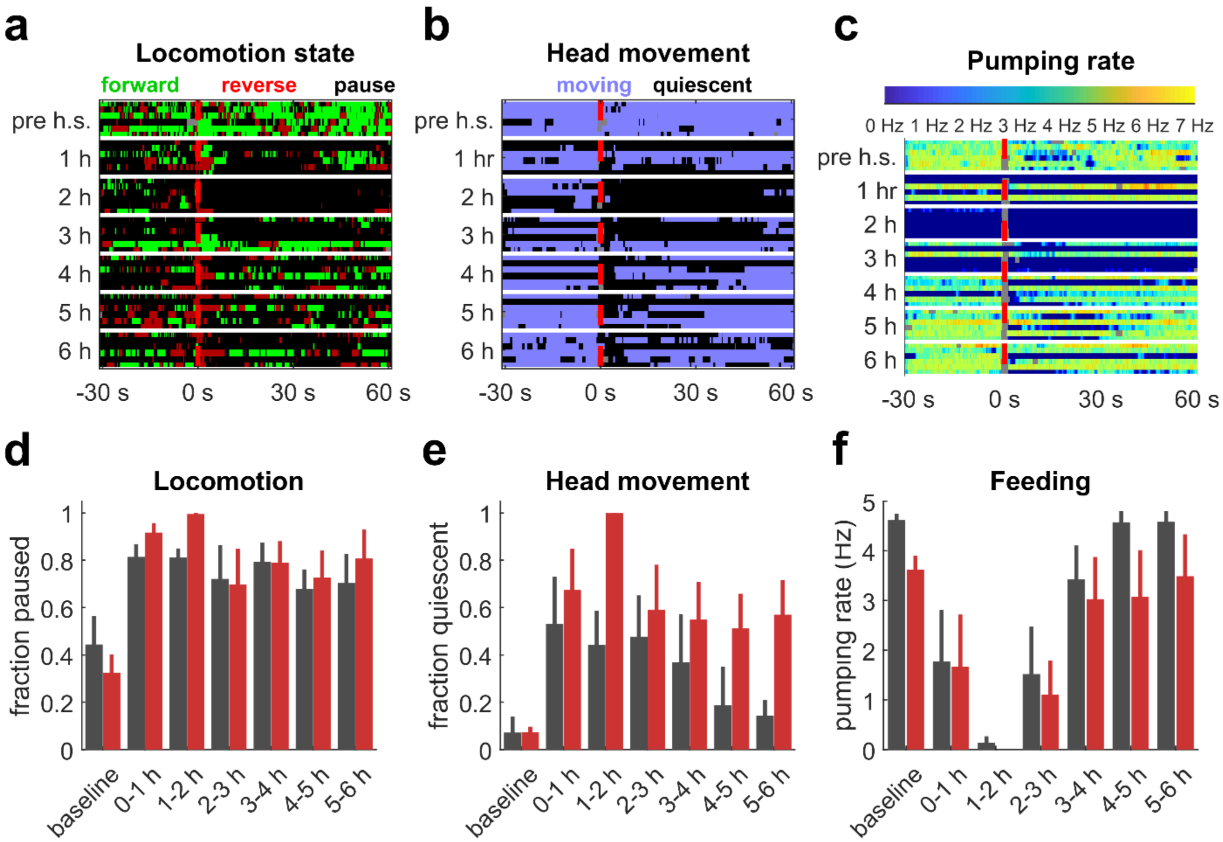


**Supplementary Figure S14. Locomotion, head movement, and feeding before and after EGF overexpression and mechanostimulus.**
**(a-c)** Heat maps showing locomotion state, head movement state, and pumping rate from 30 s before to 60 s after a 1 s, 1 kHz substrate vibration (dashed red lines) in hsp:EGF animals before a 10 min 32°C heat shock and at 1, 2, 3, 4, 5, and 6 h afterward. n = 6 animals, seven recordings each. Censored data are in gray. Data from the same animals are used in panels d-f and Fig. 6.
**(d-f)** Bar graphs showing average locomotion and head movement quiescence and pumping rate before (gray) and after (red) mechanostimulus. n = 6 animals. Error bars represent SEM.

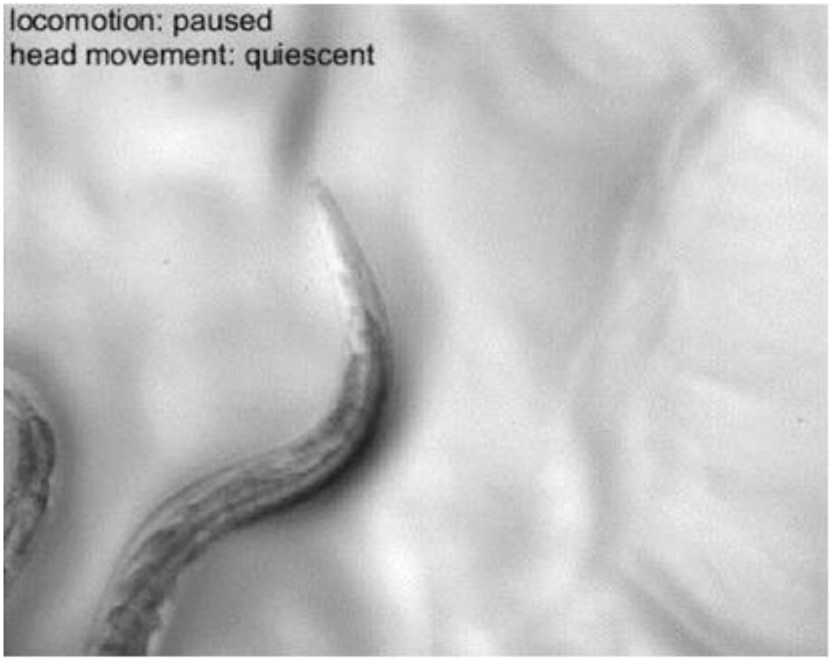


**Supplementary Video 5. Manual scoring of locomotion and head movement state.**
The recording shown here was 50 fps and 40 s in duration, but this clip has been downsampled to 5 fps.
